# The Effect of Anthropogenic Land Cover Change on Pollen-Vegetation Relationships in the American Midwest

**DOI:** 10.1101/037051

**Authors:** Ellen Kujawa, Simon Goring, Andria Dawson, Randy Calcote, Eric Grimm, Sara Hotchkiss, Elizabeth A. Lynch, Jason McLachlan, Jeannine-Marie St-Jacques, Charles Umbanhowar, Jack Williams

**Affiliations:** Nelson Institute for Environmental Studies, University of Wisconsin – Madison; Department of Geography, University of Wisconsin – Madison; Department of Statistics, University of California, Berkeley; Department of Geosciences, University of Arizona; Limnological Research Center, University of Minnesota; Illinois State Museum, Springfield, Illinois; Department of Botany, University of Wisconsin – Madison; Biology Department, Luther College; Department of Biological Sciences, University of Notre Dame; Prairie Adaptation Research Collaborative, University of Regina; Departments of Biology and Environmental Studies, St. Olaf College; Center for Climatic Research, University of Wisconsin – Madison

**Author notes:** Corresponding author: Ellen Kujawa Present Address: 550 N Park St, Madison WI 53706 USA.

## Abstract

Fossil pollen assemblages are widely used to reconstruct past vegetation community composition at time scales ranging from centuries to millennia. These reconstructions often are based on the observed relationships between the proportions of plant taxa in the source vegetation and the proportions of the corresponding pollen types in pollen assemblages collected from surface sediments. Pollen-vegetation models rely upon parameters whose values typically are assumed to be stable through time, but this assumption is largely unevaluated, due in part to the rarity of comprehensive forest data, particularly for earlier time periods.

Here we present a new dataset of early settlement-era pollen records for the upper Midwest of North America and combine it with three other pollen and forest composition datasets to assess the stability of the relationship between relative pollen composition and relative abundances of tree genera for two time periods: immediately prior to Euro-American settlement, and the late 20th Century. Over this time interval, Euro-American settlement resulted in widespread forest clearance for agriculture and logging, producing major changes to forest composition and structure and the pollen assemblages produced by these forests. These major changes provide an opportunity to test the constancy of the relationship between pollen and forest vegetation during a period of large vegetation change. Pollen-vegetation relationships are modeled, using a Generalized Linear Model, for thirteen upper Midwestern tree genera.

We find that estimates of pollen source radius for the gridded mesoscale data are 25-85 kilometers, consistent with prior studies. Pollen-vegetation relationships are significantly altered for several genera: Fagus, Betula, Tsuga, Quercus, Pinus, and Picea (p < 0.05). The use of contemporary pollen-vegetation relationships to model settlement era community composition significantly under-predicts the presence of Fagus, Betula, Tsuga, Quercus and Picea at all tree densities. Pinus is over-predicted at low relative proportions (<25%), but under-predicted at greater abundances. The divergence of pollen-vegetation relationships appears to be greatest for late-successional taxa characterized by high shade tolerance and low fire tolerance, although the statistical power is low for this analysis.

Hence, the ongoing rapid changes in land use and ecological communities associated with the Anthropocene affect not just our ability to make confident ecological forecasts for the future, but can also modify our inferences about the past. In the Anthropocene era, characterized by its rapidly changing vegetation and climates, paleoecology must move from its traditional reliance on spatial calibration datasets assumed to represent a single “present”. Instead, when possible, paleoecologists should develop calibration datasets of pollen and forest composition that are distributed across major vegetation changes in time and space. Multitemporal calibration datasets are increasingly possible given the growing length and availability of vegetation observational data and will enable paleoecologists to better understand the complex processes governing pollen-vegetation relationships and make better-informed reconstructions of past vegetation dynamics.

## Introduction

Fossil pollen records and other paleoecological proxies provide ecological insights that are otherwise temporally inaccessible, expanding knowledge of community composition, environmental change, and ecological dynamics over centuries to millennia. This understanding of long-term community dynamics can be applied to understand ecological patterns and processes in a number of ways, including reconstruction of historical ecosystems (Birks et al. 2000, Morris et al. 2014), assessing climate-driven species range expansions and contractions (Davis and Shaw 2001, Morris et al. 2014), studying the causes of abrupt change in ecological systems (Booth et al. 2012), understanding the processes that give rise to the emergence of no-analog communities (Jackson and Overpeck 2000, Williams and Jackson 2007), and determining historical baselines for restoration goals and historical patterns of disturbance (Birks 1996, Foster and Motzkin 2003).

Knowledge of the relationship between pollen composition and its source vegetation is imperative for accurate reconstruction of past vegetation composition and structure. This relationship is complicated by differences among taxa in dispersal vector, pollen productivity, and dispersibility (Cain 1939, Erdtman 1943). Relative abundance data are further complicated by the Fagerlind effect, in which the abundance of one taxon depends on the abundance of other taxa at the site (Fagerlind 1952, Davis 1963, Prentice and Webb III 1986, Davis et al. 1991).

Empirical investigations into pollen-vegetation relationships typically rely upon spatial networks of paired pollen assemblages and vegetation observations (e.g. Bradshaw and Webb III 1985, Davis et al. 1991, Calcote 1995, Parshall and Calcote 2001, Williams 2002, Williams and Jackson 2003). These studies consistently show a positive relationship between pollen and plant abundances, and indicate an effect of basin size on pollen source area, with smaller basins capturing pollen from smaller source areas than larger basins (e.g. Tauber 1965, Jacobson and Bradshaw 1981, Bradshaw and Webb 1985, Prentice 1985, Sugita 1994). Estimates of pollen source area also depend on the spatial resolution of the study (Sugita 1994) with local-scale studies tending to indicate a source area on the scale of tens of meters (e.g., 40-50% of pollen in small forest hollows coming from within 50-120 meters of the pollen source (Jackson 1990, Calcote 1995, Jackson and Kearsley 1998) Regional-scale studies with spatial resolution of 1km or larger tend to produce correspondingly larger estimates of pollen source area, on the order of 10-50 km radii or larger (Bradshaw and Webb 1985, Williams and Jackson 2003, Dawson et al. in revision).

Strategies for modeling vegetation composition from pollen data are diverse and have become increasingly sophisticated. Early methods ranged from relatively simple linear regression and ratio-based models (Davis 1963, Webb III et al. 1981, Bradshaw and Webb III 1985), to extended R-value (ERC) models, which address the Fagerlind effect (Parsons and Prentice 1981, Prentice and Webb III 1986, Prentice et al. 1987). The Prentice-Sugita family of models use dispersal kernels based on the Sutton equations (Sutton 1953) for particulate dispersal for ground-level sources and information about pollen fall speeds and productivity to estimate the pollen loadings produced by plant communities around a lake (Prentice 1985,1988, Sugita 1994). The most recent versions, LOVE and REVEALS (Sugita 2007a, 2007b), make up the Landscape Reconstruction Algorithm (Sugita et al. 2010), in which large lakes are used to estimate regional pollen rain and small lakes are used to infer local vegetation. The LRA has become widely used to reconstruct vegetation at local to continental spatial scales (Gaillard et al. 2010). Bayesian hierarchical models have been developed recently to model pollen-vegetation relationships while also quantifying key parameters such as pollen productivity and dispersibility and the uncertainty around these estimates (Dawson et al. in revision, Paciorek and McLachlan 2009).

Regardless of form, all pollen-vegetation models are calibrated using data for one time period, then applied to make inferences for another time period. This requires, however, assuming that the relationship between pollen and source vegetation composition is constant over time. The form of this assumption varies among models. Mechanistic dispersal-based models such as REVEALS assume that the parameter estimates for dispersal, productivity, and turbulence are constant over time, while models that rely upon spatial datasets of forest and pollen relative abundance must assume that these relationships between pollen and plant abundances are stable through time. This assumption of constancy has been previously noted (e.g. Sugita 1994, Parshall and Calcote 2001), and examined. For example, pollen-vegetation relationships vary according to forest type (Seppä 1998), alterations in the structure and heterogeneity of vegetation landscapes (Bunting et al. 2004),and pollen productivity (Meltsov et al. 2011, Baker et al. 2015).

In eastern North America, because the structure and composition of contemporary forests have been heavily altered by the land use and land cover change accompanying Euro-American settlement, paleoecologists have long sought to reconstruct forest composition and pollen-vegetation relationships just prior to European settlement, known as the presettlement era (e.g. Grimm 1984, Jacobson and Grimm 1986, Davis et al. 1991, Hotchkiss et al. 2007). In Minnesota, the relationships between pollen and climate have shifted significantly from the pre-settlement era to the present (St-Jacques et al. 2008a) and this can affect pollen-climate reconstructions. St-Jacques et al. (2015, 2008b) recommended that pollen-based paleoclimatic inferences should be based on pre-settlement era pollen data, rather than modern pollen-climate calibration sets, to minimize the distortion and bias produced by 19th- and 20th-century land use.

Here we build upon prior work by building novel combination of four contemporary and pre-settlement pollen and vegetation datasets, and analyzing these data to 1) quantify the relationship between the relative abundances of pollen types and tree taxa for thirteen tree genera present in the upper Midwest, 2) estimate pollen source area, and 3) assess the constancy of pollen-vegetation relationships from just at or before time of Euro-American settlement (hereafter called pre-settlement) to the present. We develop a new dataset of pre-settlement fossil pollen assemblages, drawn from the Neotoma Paleoecology Database (Grimm et al. 2013), and individual data contributors. We combine this pre-settlement pollen dataset with datasets of forest composition measured at the very beginning of the Euro-American settlement era, based on Public Land Survey System (PLSS) data (Goring et al., submitted), modern pollen data from the Whitmore et al. database (Whitmore et al. 2005), and modern vegetation composition based on the U.S. Forest Service’s Forest Inventory and Analysis (FIA) plots compiled in Goring et al. (submitted). We build a series of generalized linear models (GLMs; Rigby et al. 2005) to test whether the relationship between pollen and tree relative abundances differ among taxa (as expected, based on prior work), and between the pre-settlement era and present. Note that the pairs of pollen-vegetation datasets are not perfectly temporally aligned, e.g. the ‘modern’ pollen assemblages were collected from the 1960’s to early 2000’s while the FIA data are from 2007-2011, and the pollen data are pre-settlement while the PLS data range from pre- to early settlement. However, given the large changes in vegetation composition between the present and pre-settlement eras, the effects of these temporal mismatches should be small. To estimate pollen source area, we determine which radius for averaging vegetation composition around pollen depositional sites produces the best goodness of fit in GLMs, and compare these estimates to prior work (Calcote 1995, Williams 2002). We assess the amount of divergence between the pre-settlement era and contemporary pollen-vegetation models for individual tree genera and discuss the possible role of ecological traits such as pyrophily and shade tolerance in explaining these differences.

## Euro-American Land Use and Land Cover Change in the Upper Midwest

Euro-American land use in the upper Midwest was characterized by extensive land clearance for agriculture and forest harvesting for timber products. Prior to Euro-American settlement, forests in the upper Midwestern states of Michigan, Minnesota, and Wisconsin comprised several distinct community types, most notably the hardwood and boreal forests (north and east of the prairie-forest community border) and the prairies and oak savannas (south and west of the prairie-forest community border, Curtis (1959)). Compositional ecotones among forest types were steeper than at present (Goring et al. submitted). Southern oak savanna and prairie communities were largely maintained by frequent fires(Curtis 1959, Shea et al. 2014). By 1935 the oak savanna had essentially disappeared and had been replaced by agricultural fields (51% of the overall landscape) and pasture (ll%)(Rhemtulla et al. 2007). The overall effect of settlement in the south was the elimination of oak savannas, the near-elimination of prairie, and the replacement of these ecosystems by agricultural land and late-successional, shade-tolerant deciduous forests.

In northern Wisconsin, upland mesic pre-settlement era forests were composed of northern hardwood species (Acer spp., Betula papyrifera and B. allegheniensis, Tilia americana, and Ulmus spp.) and Tsuga canadensis; lowland forests were dominated by Larix laricina and Thuja occidentalis (Curtis 1959). In Wisconsin in 1850, approximately 84% of this northern region was forested; by 1935 the amount of forested land decreased to 56% (Rhemtulla et al. 2007). The logging of northern forests reduced conifer biomass and community diversity and led to their subsequent replacement by deciduous forest species (Goring et al. submitted, Rhemtulla et al. 2007). Northern logging resulted in declines of Pinus, while widespread slash burning and continued logging led to a decrease in fire-intolerant boreal species such as Tsuga canadensis, and an increase in early successional genera such as Betula and Populus (Rhemtulla et al. 2007). About 1.1% of the settlement era forests remained unlogged in Michigan, Minnesota, and Wisconsin as of 1995 (Frelich 1995). Anthropogenic fire suppression after the 1930’s also has altered forest composition, particularly in sandy areas with high fire-return inttervals (Heinselman 1973, Johnson 1976b, Friedman and Reich 2005). Currently, modern fire return intervals are an order of magnitude less frequent than fires before Euro-American settlement, in most areas (Cleland et al. 2004). However, fire return intervals may have increased in regions formerly occupied by Tsuga forests, where pre-settlement fire return intervals likely were >1000 years. In some regions changes have led to an increase in shade-tolerant, fire-sensitive, mostly deciduous species in regions previously dominated by fire-tolerant conifers (Drobyshev et al. 2008), and a decrease in shade-intolerant, fire-tolerant, or early-successional species in areas that previously had fire-adapted vegetation (Curtis 1959, Friedman and Reich 2005, Rhemtulla et al. 2007), however regional vegetation, and species associations vary across the region, so generalizations should be used with caution. Recent estimates suggest that 29% of modern forests in the upper Midwestern US have no historical analogue when examined at a regional (64 km^2^) scale (Goring et al. submitted).

Abandonment of farmland during the Great Depression (1929-1939) was especially common in the north, where the land was poorly suited to agriculture (Johnson 1976b, Gough 1997). Some of this land was naturally reforested or planted with seedlings in the by public work relief programs like the Civilian Conservation Corps (Johnson 1976b), which planted mostly *Pinus resinous* seedlings on cleared land (Ahlgren and Ahlgren 1983). Reforesting has not significantly affected total area of forested land in the upper Midwest over the post-Euro-American-settlement era, because of the concurrent loss of forests to the expansion of farmland in the south, but has affected local forest composition and structure (Rhemtulla et al. 2007).

## Data and Methods

### Pre-Settlement Pollen Dataset

The new pre-settlement pollen dataset presented here consists of 212 sites in Michigan, Minnesota, and Wisconsin (Figure 1a, Supplemental Table 1), and is compiled from multiple sources. Of these, 127 pre-settlement era pollen samples were obtained from the Neotoma Paleoecology Database, which is a community-curated, open-source collection of paleoecological data, (Grimm et al. 2013, http://www.neotomadb.org). We obtained data using the neotoma package for R (Goring et al. 2015), searching for all sedimentary cores within the upper Midwest that had samples within the most recent 2000 years, and particularly within the last 500 years. In addition to these sites, we included 21 presettlement samples from a dataset gathered by Ed Cushing, that has been previously published, expanded and used elsewhere (Jacobson and Grimm 1986, St-Jacques et al. 2008a, 2008b, 2015). To these we added pre-settlement samples that have been recently published and/or contributed by researchers. We uploaded some of these to Neotoma (Supplementary Table 1; all records with Neotoma IDs >10,000), while other presettlement samples are part of pollen datasets that are being prepared for submission to Neotoma (Supplementary Table 1; all records with “CLH” or “CALPRE” in the ID column).

**Table 1.**
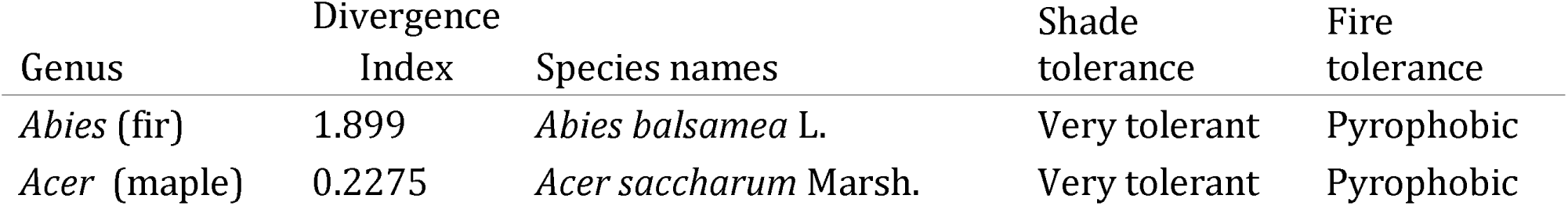
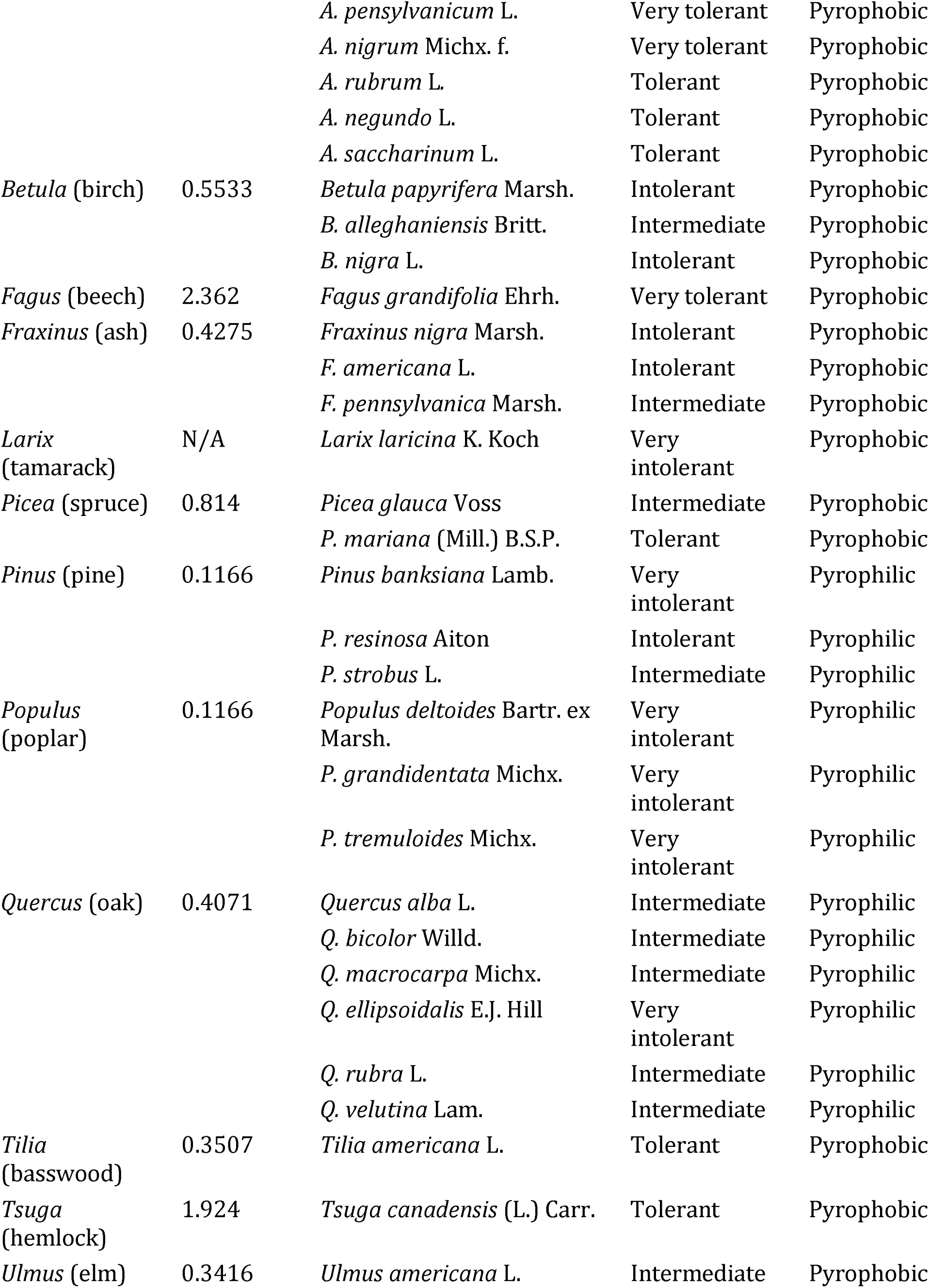
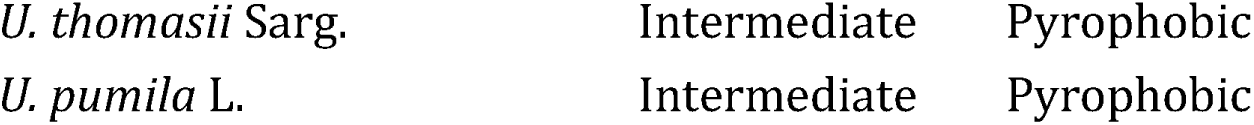
An index of model divergence. The index value is the mean of absolute differences between modern and pre-settlement predictions divided by the pollen proportion expected at the pre-settlement era. Shade tolerance information is sourced from Gower (1996), with the exception of Acer pensylvanicum (Gabriel and Walters 1990), Fraxinus pennsylvanica (Kennedy 1990), Betula nigra (Grelen 1990), Quercus ellipsoidalis (Coladonato 1993), and Quercus bicolor (Snyder 1992). Fire tolerance information is from Thomas-Van Gundy et al. (2015). Non-native tree species are excluded from this analysis.

**Figure 1.**
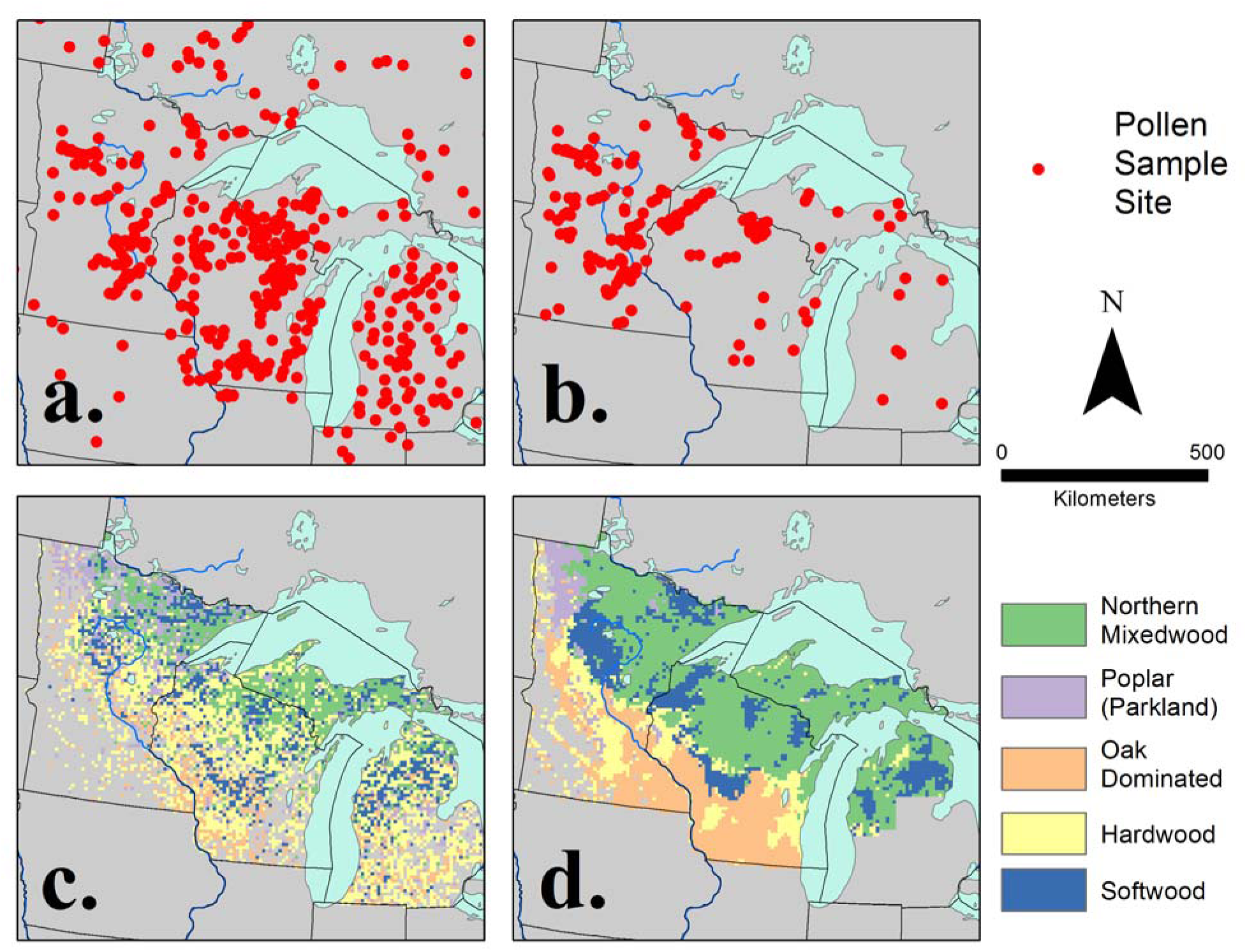
Map of pollen sites in Michigan, Minnesota, and Wisconsin (a, b) available for the late 20th century (a: modern era) and just prior to major Euro-American settlement and land clearance (b: pre-settlement era and modern). Forest types are classified using cluster analysis from the FIA (c) and PLSS (d) forest data (Goring et al. submitted). As discussed in Goring et al. (submitted) and elsewhere, Euro-American land use produced major shifts in forest composition and heterogeneity.

Most pre-settlement samples obtained from the Neotoma Paleoecology Database (including previously archived records and the ones newly added here) were extracted from longer fossil pollen records from cores and stratigraphic sections. For each of these records, the pre-settlement horizon had to be identified. Fortunately, the Euro-American settlement horizon is a well established biostratigraphic feature in eastern North America, largely identified by significant increases in *Ambrosia* and other associated “weedy” taxa such as Rumex (McAndrews 1988). We used an expert elicitation exercise to identify the settlement horizon. Expert elicitation uses expert knowledge to make estimates from (often) noisy data (Bond et al. 2012). Here four palynologists with experience in the upper Midwest were asked to identify the uppermost pollen samples prior to the settlement horizon (Dawson et al. in revision). Sites were discarded if sampling resolution was too low, defined here as a gap of >150 years between the age of the pre-settlement pollen sample and the start of settlement period. These pre-settlement data were supplemented by 48 mostly unpublished records from across northern Wisconsin (Supplemental Table 1), contributed by Calcote, Lynch, and Hotchkiss (Hotchkiss et al. 2007, Lynch et al. 2014). For these samples, pre-settlement samples were identified as the last sample before an increase in *Ambrosia* pollen percentage above background, and the decline of white pine (due to logging) which often occurs just before the *Ambrosia* rise. These samples were generally within 100 years before 1850. All pollen counts from this pre-settlement-era dataset have been made publicly available as part of the GitHub repository for this paper http://githb.com/PalEON-Proiect/Pollen Vegetation-Kuiawa and as supplementary material.

### Pre-Settlement Vegetation: The Public Land Survey System

The Public Land Survey System (PLSS) began as a result of the Land Ordinance Act of 1785 and the Northwest Ordinance Act of 1787 (Carstensen 1988). Surveyors used a gridded system of six-mile-square areas, termed “townships”, which were subsequently divided into 36 one-mile-square sections. At the corners and midpoints of these subplots, and at meander corners (locations where section lines intersected navigable bodies of water), surveyors recorded the species and diameter at breast height (DBH) of the nearest witness trees and their cardinal direction and distance from the point (Schulte and Mladenoff 2001). Roughly 69% of the 48 continental United States is continuously covered by this rectangle-based survey, and an additional 9% is covered intermittently (Johnson 1976a).

PLSS data represent one of the broadest and most detailed records of pre-settlement era vegetation. As such, they have been used to address a wide variety of ecological questions, and to reconstruct the community composition of pre-settlement forests (Anderson and Anderson 1975, Kline and Cottam 1979), quantify land use changes (Radeloff et al. 1999, Leahy and Pregitzer 2003), and establish restoration baselines (Fritschle 2012). Accuracy of PLSS data is affected by imprecise species identification and surveyor bias toward easily accessible, larger, or more valuable trees, particularly at fine spatial scale (Schulte and Mladenoff 2001, Fagin and Hoagland 2011, Liu et al. 2011). Nonetheless, these witness tree records provide the most complete assessment of tree composition in the western and central United States just before time of Euro-American settlement.

The data used in this analysis were sourced from a standardized dataset presented in Goring et al. (submitted). Goring et al. built on prior work, (e.g. Kronenfield and Wang 2007, Rhemtulla et al. 2009, Liu et al. 2011, Hanberry 2013) and includes novel correction factors to remedy issues of surveyor bias and sampling geometry in the original records, creating a dataset of gridded (8x8km) estimates of species composition (Figure 1d; used in our analysis), basal area, stem density and biomass. The analysis calculates stem density as a function of the distances of the two trees closest to the plot center. Corrections use a modified Morisita estimator, where stem density is calculated at the individual point, and then aggregated to the 8x8km grid cell. Estimating at the point level permits the application of point-specific corrections, to account for the fact that, for example, plot geometry varies between section and quarter-section points. The PLS data then provides regional and continuous coverage of vegetation, at, or immediately prior to major land use conversion by Euro-American settlers across Minnesota, Wisconsin, the Upper Peninsula of Michigan and the upper third of the Lower Peninsula (Goring et al. submitted).

### Modern Vegetation: Forest Inventory and Analysis

The research branch of the United States Forest Service (USFS) was established in 1928 as a result of the McSweeney-McNary Forest Research (Shaw 2008, USFS 2014). The McSweeney-McNary act specifically instructed the USFS to maintain a comprehensive census of the United States’ timber resources (Shaw 2008), and the USFS began Forest Surveys in 1930(USFS 2015). The Forest Survey program has since been renamed the Forest Inventory and Analysis National Program (FIA); currently, the 48 contiguous U.S. states and parts of southern Alaska are surveyed annually (USFS 2014).

FIA data have been used in climate change research (Domke et al. 2014, MacLean et al. 2014), biodiversity conservation (Hanberry 2013, Randolph et al. 2013, Belote and Aplet 2014), and timber analysis (Domke et al. 2014). FIA data are widely applicable to ecological study because they represent a relatively complete picture of the vegetation (particularly arboreal taxa) of the United States: approximately 820 million acres are included in FIA plots and over three hundred thousand plots are subject to comprehensive vegetation surveys annually (USFS 2014).

FIA data in this study are from 2007-2011, and were prepared as in Goring et al. (submitted). Plot level data were aggregated to 8x8 km cells. The number of FIA plots per cell varies, but across the region there is a median of 3 FIA plots per cell (Goring et al., submitted). Within each cell, all trees with diameters >8” were summed to generate an estimate of percent composition for that cell. This diameter cutoff was chosen for consistency with the PLSS data and estimates of minimum tree size used by Public Land Surveyors (Goring et al. submitted).

Comparisons between FIA and PLSS forest data are potentially made more difficult by differences in methods and scales between datasets. These differences are, to some degree, addressed by the common scale of aggregation (8km) used for both datasets. To further address this issue, Goring et al. (submitted) performed sensitivity analysis to understand the potential effects of differences in sampling methods and plot resolution between PLSS and FIA era data on the apparent changes to the landscape following Euro-American settlement, known as the “mixed pixel” problem (Kronenfield et al. 2010). Results showed that differences in composition are not related to the intensity of sampling within plots for the FIA or PLSS, and so, at this scale, are likely to represent true differences, rather than artifacts of differential sampling systems between the PLSS and FIA datasets (Goring et al. submitted).

### Data Preparation

Data from Michigan, Minnesota, and Wisconsin were used for this analysis. Data from a set of thirteen arboreal taxa were present in all four datasets: *Abies, Acer, Betula, Fagus, Fraxinus, Larix, Tilia, Picea, Pinus, Populus, Quercus, Tsuga, and Ulmus*. We considered including a “other” category to represent other tree taxa and non-arboreal taxa, but decided that this category would be incomparable among datasets and time periods, because of differences in sampling method. To ensure consistency among all four datasets, percentages reported in all datasets represent proportions of the summed values of only these thirteen genera. The use of genera (Table 2) is necessary, given taxonomic uncertainty in both pollen and PLSS datasets. Because pollen is identified based on morphological features, and pollen morphology is strongly conserved, it is often difficult to differentiate pollen types to the species level (although it is possible, e.g. for some *Pinus* sp., *Picea*, and other taxa; Hansen and Cushing 1973, Lindbladh et al. 2002,2003, May and Lacourse 2012, Mander and Punyasena 2014). Similarly, PLSS trees were identified to the species level by surveyors, but often recorded using common names that were locally idiosyncratic or are ambiguous, resulting in uncertainty below the genus level when trees are identified using terms such as “Pine” or “Oak”. Although methods exist to differentiate PLSS data to the species level using information about local edaphic conditions (Mladenoff et al. 2002), here we opted to work with both the PLSS and pollen data at the genus level.

All datasets were projected to Albers equal-area conic projection using the R package rgdal (Bivand et al. 2013). For each pollen site, we calculated proportional vegetation for each of the thirteen taxa based on the average of all vegetation grid cells within a distance radius. This distance radius varied in 10 km intervals from 5 to 200 km, to encompass potential source areas found in other regional-scale pollen-vegetation studies (Bradshaw and Webb, 1985; Dawson et al. in revision).

### Data Analysis

To model the relationships between pollen and vegetation and test hypotheses about the stability of pollen-vegetation relationships, we used the gamlss package (Rigby and Stasinopoulos 2005) to build three successively more complex generalized linear models (GLMs). The gamlss package is typically used to build generalized additive models (GAMs), flexible multivariate regression models that incorporate nonlinear predictive models (Hastie and Tibshirani 1986); however, gamlss may also be used to create GLMs, by omitting smooth splines, and contains more options for distributions than other GLM packages. We used a zero-inflated beta distribution, which is well-suited to proportional data with many zeros like ours (tree genera that are not present at any site receive a zero for that site) (Ospina and Ferrari 2010) We modeled pollen proportion as a function of vegetation proportion, with additional covariates including genus and data collection period (pre-settlement era or modern) as the predictor variables.

We tested models of increasing complexity to assess the significance of taxa and time as covariates. In Model 1, we estimate relative pollen composition as a function of vegetation, assuming that all genera and time periods behave identically. The second model, Model 2, includes taxon as a covariate of vegetation, so that the relationship is allowed to vary by taxonomic group, while the third model, Model 3, also includes an era covariate representing the time period of the vegetation dataset (either pre-settlement or the present era). Model 3 allows us to explicitly test whether there is a statistical difference in pollen-vegetation relationships for each genus between the time of Euro-American settlement and the present. We used GAIC (Generalized Aikaike’s Information Criterion, Rigby and Stasinopoulos 2005) to assess goodness of fit for each model and to determine the optimum radius (a proxy for pollen source area), by choosing the model and distance radius with the lowest AIC value. Vegetation is weighted using an inverse square distance weighting. Preliminary analysis indicated that this weighting provided the best fit for our models.

To quantify the magnitude and importance of the difference between pollen-vegetation models based on pre-settlement data versus models based on modern data, we calculated a divergence index for each genus, by taking the mean of absolute differences between modern and pre-settlement predictions, divided by the pollen proportion expected at the pre-settlement era. This standardization corrects for the expectation that more abundant taxa should have larger differences in their relative abundances. To assess whether shifts in disturbance history or successional stage might be affecting the observed shifts in pollen-vegetation relationships, we compared the divergence index to fire- and shade-tolerance levels for that genus (Table 1). We applied a Bonferroni correction (using the R function ‘p.adjust’ to the *p*-values returned from the ANOVA comparisons.

The ggplot2 (Wickham, 2005) and gridExtra (Auguie 2015) packages were used to construct figures in R (R Core Team 2014).

## Results

### Pollen-Vegetation Relationships

The most complex model (Model 3), which models pollen proportion as a function of vegetation proportion, taxon and era provides the best fit based on the AIC (Figure 2). For each of the three models considered, the lowest AIC (indicating best goodness-of-fit) corresponded with vegetation data that were averaged using distance radii on the order of tens of kilometers. Model 1 had the lowest AIC at a distance of km, and Models 2 and 3 had the lowest AIC at a distance of km respectively (Figure 2). This in turn implies that the source area for these lakes or bogs is on the order of km. Spatial smoothing in the original 8km dataset and the search radius used here reduces the variability of vegetation proportions, so that few pollen sites have 0% proportions for particular tree taxa (e.g., Figure 2e), while also reducing the highest proportions for certain taxa (e.g., Figure 2b and c). At very large radii (>100 km) the estimates of vegetation revert to some mean estimate for each taxon, making a pollen-vegetation relationship difficult to recover, resulting in higher overall AICs. Model improvement in RMSE for Model 3 is approximately 25% from the single-grid-cell radius (RMSE_cell_ = 8.9%) to the model employing an 85km radius (RMSE_85km_ = 6.6%).

**Figure 2.**
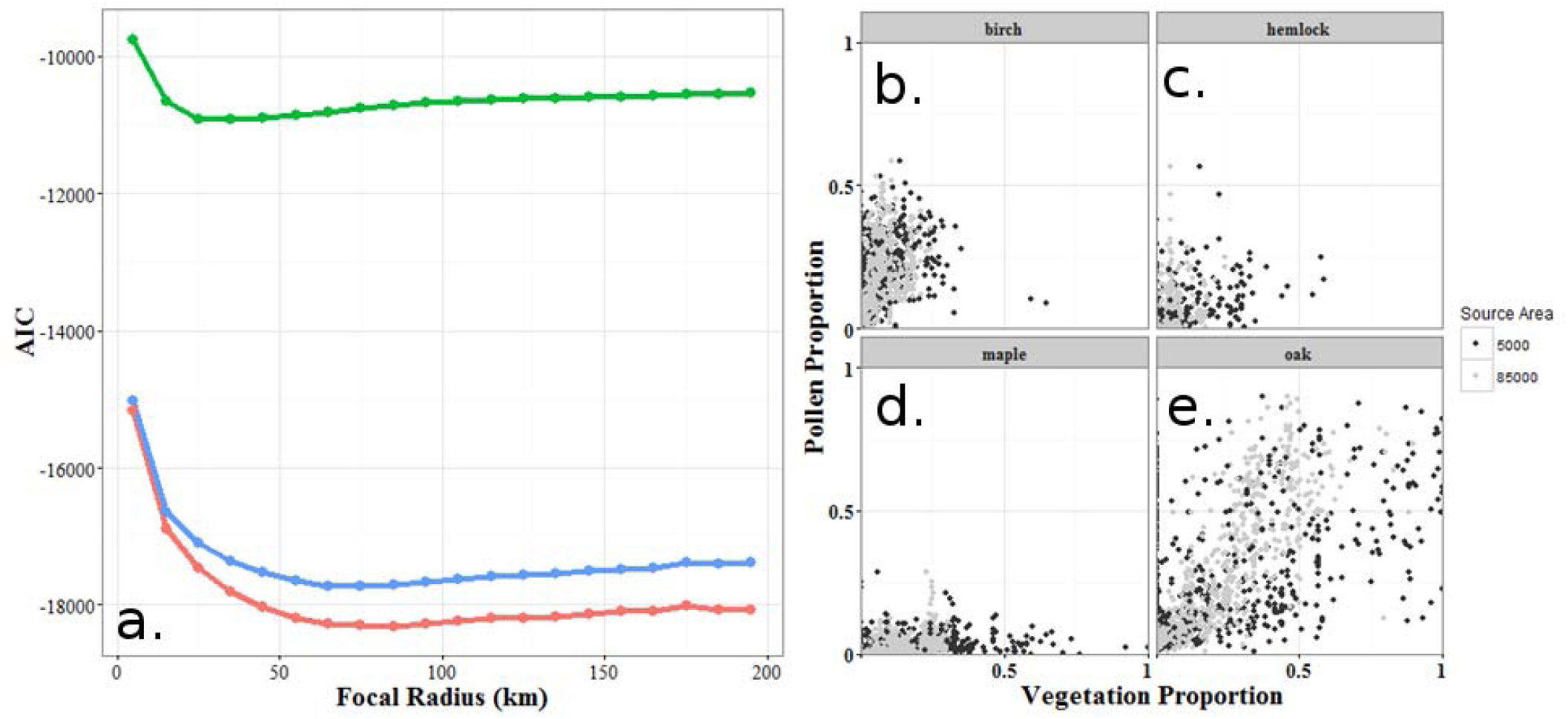
Aikaike’s Information Criterion (AIC) values for three models at10-km intervals from 5-200 km (panel a). Model 1 (green) has the highest goodness of fit (lowest AIC) at a distance of km; models 2 (blue) and 3 (red) have the highest goodness of fit (lowest AIC) at km respectively. To illustrate how the relationship between pollen and vegetation proportions is improved by increasing vegetation radius (and, by implication, a closer fit to pollen source area), panels b-e show the pollen-vegetation relationship for four selected genera (b: birch, c: hemlock, d: maple, e: oak) in the focal grid cell (64km^2^) and smoothed to a radius of 85km.

Model 1, a very simple model that directly relates pollen to vegetation, regardless of genus or era, indicates that the proportion of a tree genus on a landscape is a significant predictor of its proportional composition in pollen assemblages (p < 0.001), confirming that pollen composition is an indicator of vegetation composition. Model 2 shows that these pollen-vegetation relationships vary widely among genera (p ≤ 0.001), as is well-known (Davis 1963). *Betula, Pinus*, and *Quercus* are the most prolific pollen producers, with the steepest curves (Figure 3), while others including *Tsuga, Ulmus, and Larix* (not shown) are underrepresented genera that rarely reach high pollen proportions. The representation of *Larix* in the pollen record is complicated by its affinity for wetland environments. Hence, *Larix* is likely to be over-represented in the vegetation immediately adjacent to pollen sites in or near wetlands or bogs, but under-represented in the uplands between pollen sites. As a result, its relative proportion in the pollen record is more likely an indication of adjacent wetlands than of total landscape coverage around a sample site. For this reason, we include *Larix* in our models, but exclude it from our figures.

**Figure 3.**
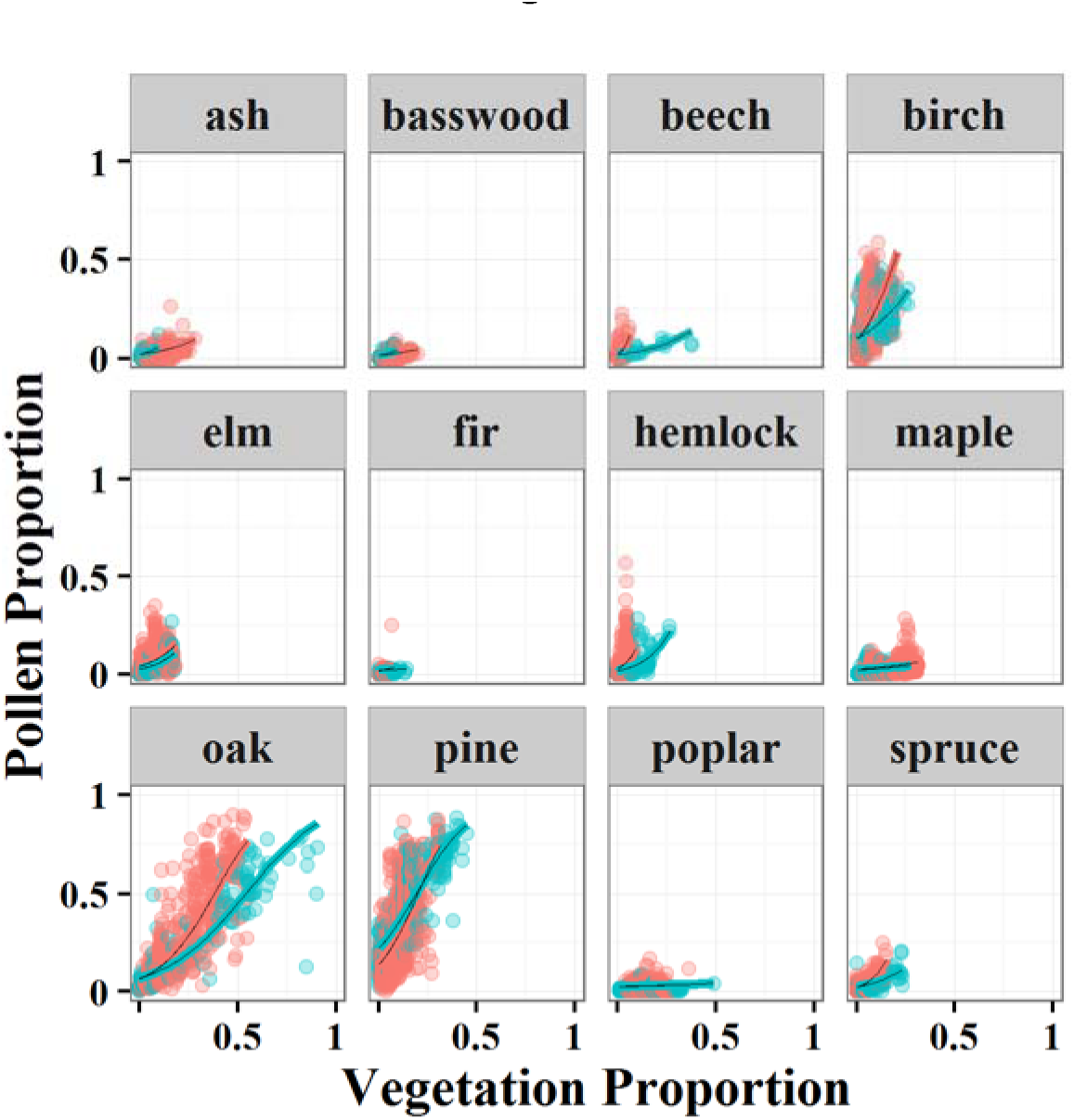
A scatterplot of pollen and vegetation proportions, using Model 3: pollen proportion as a function of vegetation proportion, genus, and period. Each data point corresponds to the proportion of total pollen per genus at a single pollen site at each sampling period, as a function of the proportion of total vegetation of the same genus and sampling period, averaged for all vegetation grid cells within an 85 km radius of the pollen site. Beech, birch, hemlock, oak, pine and spruce each have a statistically significant difference in the pollen-vegetation relationship for that genus between pre-settlement era (blue) and present (red).

In Model 3, the relationships between pollen and vegetation composition change significantly several genera from the pre-settlement era to the present: *Fagus, Betula, Quercus, Pinus, Tsuga, and Picea*. Nearly all these genera produce lower proportions of pollen relative to vegetation during the pre-settlement era than at present. The one exception is Pinus, which shows statistically significant different relationships for the two periods, but the two models cross, with Pinus over-predicted on the pre-settlement era landscape at low densities (<25%) and under-predicted at higher densities. Many of these shifts may be driven by changing vegetation proportions between pre-settlement and modern eras; for example, *Fagus* reached regional vegetation proportions of almost 50% in the pre-settlement era, but is rarely seen in proportions above 10% in modern forests. Similar declines in forested cover are seen for *Tsuga* and *Fraxinus* and, to a smaller extent, *Quercus*.

These altered pollen-vegetation relationships are also likely driven by broad-scale changes in forest composition across the region and shifts in co-occurrence between high and low pollen producers. Because pollen percentages are expressed as a relative sum, increases in the proportion of one genus can drive down the relative pollen proportions of other genera, particularly if the first genus is overrepresented in the pollen record and the other genera are underrepresented (Fagerlind 1952, Prentice and Webb III 1986). For example, the proportions of *Abies* pollen as a function of *Abies* vegetation increase strongly from the presettlement era to the modern, indicating that Abies is less under-represented in modern pollen assemblages than in the pre-settlement era. This shift in *Abies* may be caused by changes in *Pinus* vegetation proportions (a notoriously prolific pollen producer) and a weakened association between *Pinus* and *Abies* in modern forests. Correlations between *Pinus* and *Abies* vegetation decline from r_pre_settlement_ = 0.59 to r_modern_ = 0.18, indicating that rates of co-occurence between Abies and the heavy pollen producer, Pinus, are lower in modern forests than in pre-settlement era forests.

*Fagus* is also under-represented in the pre-settlement era. Within the vegetation data, *Fagus* shifts from being more strongly correlated with Quercus (r_pre-settiement_ = 0.27, r_modern_ — 0.04) to more strongly correlated with *Betula* (r_Pre—settiement_ — 0.01, r_modern_ — 0. 56). Both *Fagus* and *Quercus* are heavy pollen producers, but because the proportions of Fagus are so limited in the modern vegetation it is difficult to know how much of this shift in relationship is driven by several outliers, or by true shifts in pollen-vegetation representivity.

*Fraxinus*, which is more strongly under-represented in modern samples than in presettlement samples, shows greater correlation to *Betula* and *Pinus* in modern forests, but also much stronger gains in correlation to weak pollen producers including *Acer* (the correlation between *Fraxinus* and *Acer* shifts from 0.17 to 0.61 between the pre-settlement era and present). Hence, its shift may be due not as much to increasing co-occurrence with heavy producers, but with weak producers.

The divergences between the pre-settlement and modern pollen-vegetation models varies among genera (Table 1) and are highest for *Fagas, Tsuga*, and *Abies* and lowest for *Populas, Pinas*, and *Acer*. Correlation of these divergence values to shade- and fire-tolerance characteristics is suggestive (Table 1, Figure 6) but the statistical power of this analysis is low due to the small number of taxa (12; *Larix* omitted due to known overrepresentation of pollen in wetland settings and underrepresentation elsewhere). With these caveats in mind, suggestive patterns do appear. Shade-tolerant and very tolerant taxa such as *Fagus* and *Tsuga* have stronger divergence of pollen-vegetation relationships than intolerant taxa (Table 2, p=0.012, corrected). We also see a possible signal of fire tolerance in the results, although without statistical significance, with pyrophilic species showing lower divergence than intolerant species (p=0.16). Overall, tree genera associated with mature forest show much higher rates of divergence between models than early successional taxa such as pine, poplar and maple (although successional stage varies among maple species). However, given the sample size (n=12), this analysis is more suggestive than conclusive.

**Figure 5.**
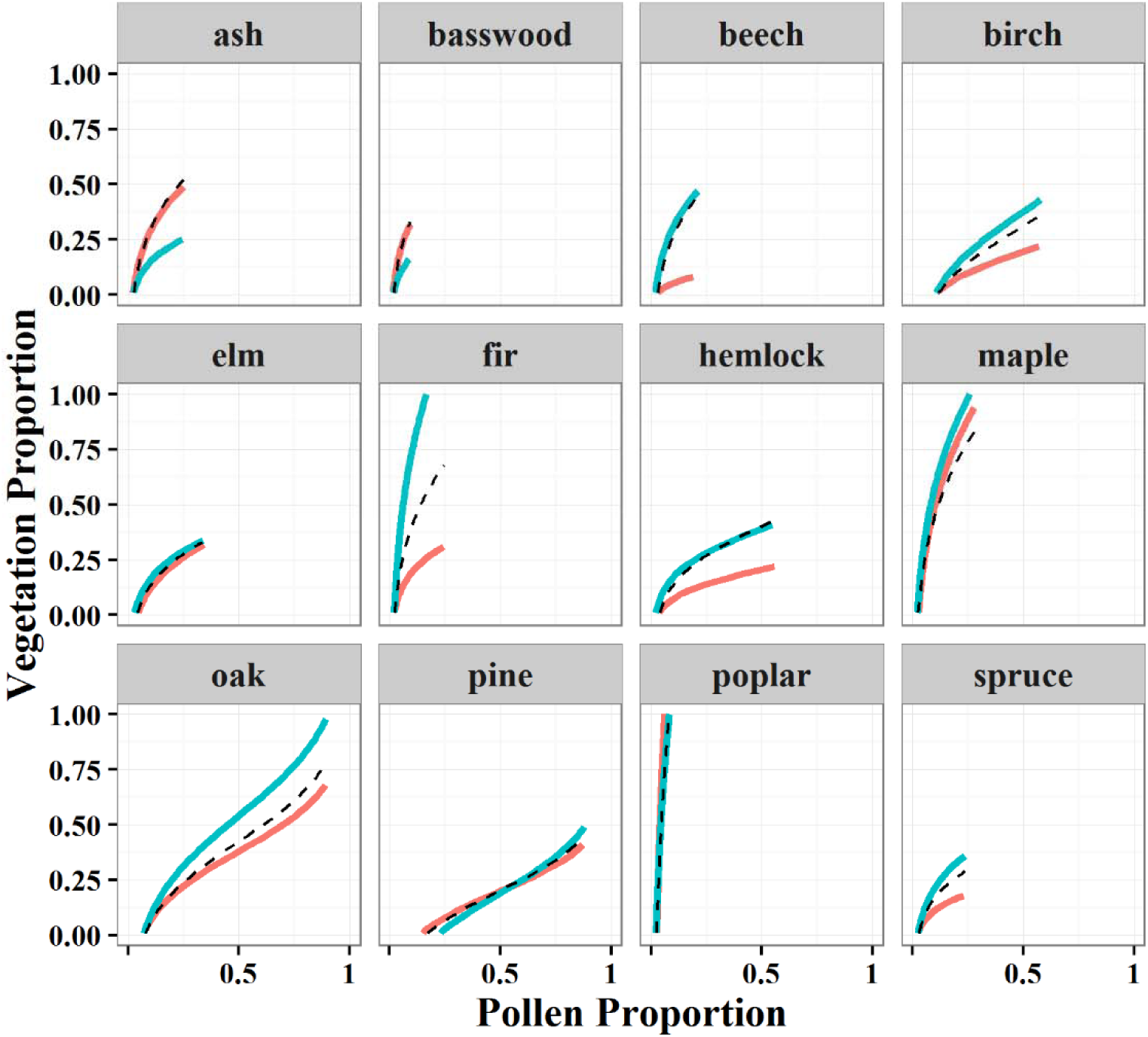
The inverted form of the pollen-vegetation relationship, in which vegetation proportion is expressed as a function of pollen proportion between pre-settlement (blue) and modern (red) eras using Model 3. Model 2 estimates are superimposed on the plot (black dashed line) for comparison. Where the pre-settlement (blue) line is above the red line, application of the modern pollen-vegetation relationship to fossil pollen records would lead to an underestimate of the relative abundance ofthat genus. The converse is true when the modern line (red) is higher than pre-settlement estimates (blue).

**Figure 6.**
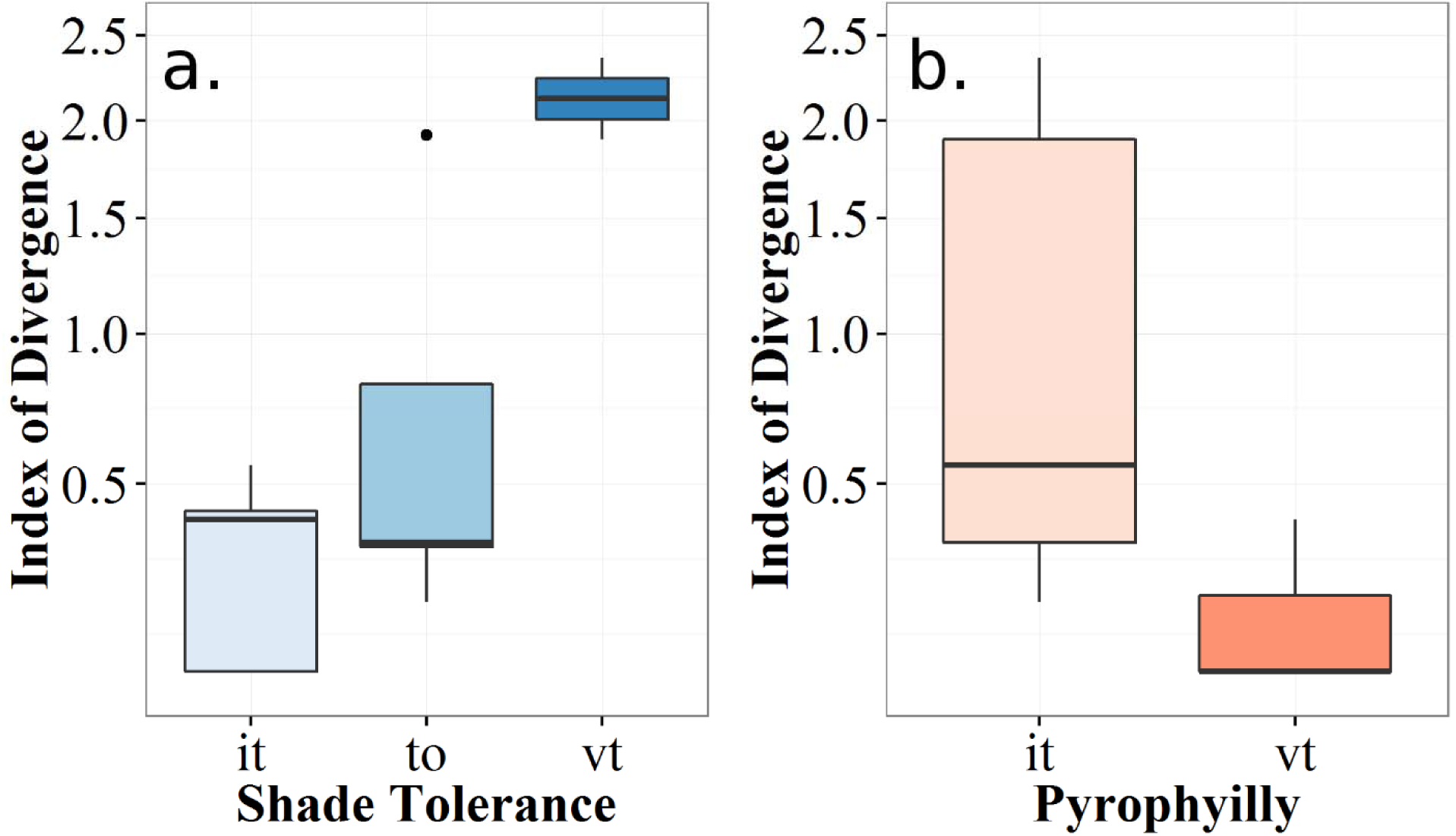
Index of divergence by tolerance for shade and fire. Each panel represents the mean (solid black bar), the 1^st^ and 3^rd^ quartiles (box limits) and the 95^th^ percentiles for the Index values within the class. Clear trends of increasing divergence are visible for shade tolerance (a) as tolerance increases from intolerant (it), to tolerant (to) and very tolerant (vt) species, significant at p=0.01 using Bonferroni correction, (b) Pyrophilly decreases divergence, indicating that species that are tolerant of fire (or that exist in fire prone landscapes; vt) have lower divergence than species that are intolerant of fire (it). These results suggest a trend, but lack significance when corrected using Bonferroni correction (p=0.16).

## Discussion

Our work confirms our understanding that pollen and vegetation taxa compositions are highly correlated, and that these relationships differ among genera (Davis 1963, Heide and Bradshaw 1982, Prentice and Webb III 1986, Webb III 1993)Our analysis also indicates that the relationships between pollen and vegetation for some genera have shifted substantially over time. Based on these findings, reconstructions of past forest composition based on modern pollen and vegetation calibration data significantly (p <0.05) underpredict the presence of some genera (*Abies, Betula, Fagus, Quercus, Tsuga;* Figure 3). These shifts in pollen-vegetation relationships are likely a response to anthropogenic land use, recent climate change, and associated changes in forest community composition.

### Estimating Pollen Source Radius and Area: Comparisons to Prior Work

Our estimates of pollen source radius (Figure 2) are consistent with prior studies. Estimates of pollen source area vary by the area of the lake basin (e.g., Calcote 1995, Sugita 2007a, vegetation structure (Bunting et al. 2004) and by the spatial resolution of the study. Our estimates of pollen source area (25-85 km radius, Figure 2) are bracketed by the source area estimates developed by Sugita Sugita 2007a, 2007b for large lakes (>100 ha) and smaller lakes. Sugita (2007a) suggested that large lakes were suitable for reconstructing regional vegetation composition at scales of 104 to 105 km^2^, (equivalent to circles with radii of 56 to 178 km). For smaller lakes, model simulations by Sugita (2007b) suggested pollen source radii on the order of 0.7 to 3.7 km.

Studies based on vegetational observation datasets with a spatial resolution of >1km and networks of moderately sized lakes (5-250 ha) located tens to hundreds of kilometers away from one other tend to find evidence for relatively large pollen source radii. Williams and Jackson 2003 found that search window half-widths of 25-75 km provided the best goodness of fit, and Bradshaw and Webb 1985 suggested that some genera accumulate significant pollen from more than 30 km away from a basin (e.g. Pinus and Quercus). Dawson et al. (in revision) estimated 50% capture radii (the maximum distance traveled by 50% of pollen sourced from a grid cell) on the order of 60 to 312 km. In contrast, modeling and empirical studies of pollen source area from forest hollows and smaller lakes tend to report smaller source areas, ranging from a few 10s of meters to a few kilometers (Calcote 1995, Jackson and Kearsley 1998, Sugita 2007b).

### Implications and Causes of Changes in Pollen Vegetation Relationships

These observed changes in pollen-vegetation relationships are non-trivial. Bunting et al. (2004) show through simulation that vegetation structure on the landscape can affect the apparent vegetation source area. Thus, the significant shift in vegetation structure and composition across the settlement boundary is likely to strongly affect pollen representivity, although here we see that it does not change the apparent source area of the optimal relationship.

The significant changes in the observed and modeled pollen-vegetation relationships between the pre-settlement era and late 20th Century (Figures 3, 4) are directly attributable to changes in vegetation composition and structure caused by human agency. Logging removed genera that were valuable timber (*Quercus, Pinus*), while the elimination of the natural predators of deer increased browsing and decreased the ability of some northern mesic and boreal species to establish (*Tsuga, Fagus*). Homogenization (Schulte et al. 2007, Rhemtulla et al. 2009) and the rise of novel forest communities (Goring et al., submitted) was also the result of human agency, resulting from regeneration following forest clearance, and much higher prevalence of pioneer species within forest communities. This homogenization also contributes to the overall “compression” of the x-axis apparent in Figure 2: *i.e.*, each plant genus has a more reduced range of relative composition in the late 20th century than it did during the pre-settlement era. Because the compositional differences among vegetation formations is lower now (Goring et al. submitted), locations that were once dominated by one or two genera are now frequently comprised of a much more diverse genus assemblage. Note that if differences in sampling design between the FIA and PLS data were the cause, one would expect to see higher heterogeneity in the contemporary data, i.e. the opposite of the observed pattern. This is because the FIA data are sampled at individual plots while the PLS data represent 2-4 trees at points spaced ∼1 mile apart, so the FIA data should be expected to be influenced more by local heterogeneity than the PLS data. Changes in vegetation heterogeneity and structure may affect pollen productivity and transport (Bunting et al. 2004), and so may be one of the causes of the changes in these pollen-vegetation relationships.

The effect of anthropogenic changes in vegetation composition and heterogeneity is magnified by the use of relative abundance data. Because the proportions of all genera at a site must sum to 1, the reduction of prolific pollen producers such as Pinus shifts the pollen-vegetation relationships for the remaining genera, and increases the apparent proportion of other genera on the landscape (Figure 4). Matthias and Giesecke (2014) show strong linear relationships between pollen accumulation rates and forest biomass in northeastern Germany. Much of the modern pollen data within the North American Modern Pollen Database (Whitmore et al. 2005) has no secondary information with which to estimate PAR, but the increasing ability of age-depth models to resolve uncertainty, and the more frequent use (and reporting) of exotic markers in pollen analysis mean that this may be a fruitful avenue for future research at broader regional scales.

Other factors may contribute to the apparent instability of pollen-vegetation relationships as well. Pollen productivity can vary from year to year, for example, at Arctic tree line, interannual variations in the pollen productivity of Picea and Pinus correlates to mean July temperatures of the previous year (Huusko and Hicks 2009). Other changes in forest structure and openness or meteorological conditions may also have contributed to the observed changes in pollen-vegetation relationships, by altering the dispersal of pollen grains. However, we believe that these factors are of secondary importance relative to the widespread changes in forest composition produced by anthropogenic land use and the consequent changes in the relative proportions of overrepresented and underrepresented pollen types.

Differences in vegetation sampling design between the two eras could, in principle, cause some of these observed changes, but we view this possibility as unlikely. As noted above, the observed changes in vegetation heterogeneity are opposite to those expected given sampling design. Goring (submitted) tested for the effects of sampling design by testing whether the degree of community dissimilarity between the 8 km grid cells in the FIA and PLS datasets was influenced by the number of FIA plots per grid cell, and reported only a minor effect. Hence, we believe that the findings reported here are robust to differences in sampling design between the PLS and FIA data.

Pollen-vegetation relationships can also vary spatially. For example, *Carya, Quercus*, and *Ulmus* have similar pollen-vegetation relationships in both the southeast and upper Midwestern United States, but *Betula, Fraxinus, Juglans*, and *Pinus* do not (Delcourt et al. 1983). This is due in part to the different genera represented in these regions (pollen types often can only be identified to genus level; different species are present in both regions, and these species have different dispersal characteristics). Dawson et al. (in revision) report that estimates of pollen productivity and dispersal obtained by a Bayesian hierarchical model (STEPPS) fitted to the upper Midwest are in closest agreement to experimentally measured values of pollen productivity and dispersal for genera also from the upper Midwest, and do not agree well with experimental values from Europe. This geographic variability suggests a need for replication of these analyses in regions outside the upper Midwest.

Other research has begun to explore how human land use may affect the information that can be retrieved from fossil pollen record, with an emphasis on pollen-based inferences of past climates. St-Jacques et al. (2008, 2015) suggest that a pollen-climate calibration set from northern Minnesota based on pre-settlement era relationships is more accurate for reconstructing climate of the past millennium than a modern pollen-climate calibration set. Their model improvement was attributed to bias in the modern pollen record due to postsettlement increases in *Ambrosia*, Chenopodiaceae, and Poaceae and decreases in arboreal genera, particularly Pinus and Quercus. Li et al. (2014) argued that pollen-based climatic reconstructions in China for the last 1,100 years were heavily biased by human land use.

### Paleoecology in the Anthropocene: New Challenges, New Opportunities

More broadly, the changing pollen-vegetation relationships documented here are part of a more general challenge of studying and understanding a rapidly changing world. Shifting baselines are a central characteristic of the Anthropocene (Jackson et al. 2001). Rates of change are accelerating, as the human imprint on ecosystems and the physical environment continues to grow (Steffen et al. 2004). Often, we use the world around us as an explicit or implicit reference point, when projecting to the future or to the past (Williams and Jackson 2007). In paleoecology and paleoclimatology, modern calibration datasets have been the backbone of quantitative research for decades (e.g. Imbrie and Kipp 1971b, Delcourt et al. 1984, Whitmore étal. 2005, Williams étal. 2006). These calibration datasets are spatially extensive but typically assumed to represent a single time period, the “present” or “modern”. However, many samples used in calibration datasets were collected decades ago (Whitmore et al. 2005). Statistical transfer functions are calibrated against these modern datasets, then applied to fossil assemblages to reconstruct past climates (Viau et al. 2002), vegetation composition and structure (Williams 2002), or other variables of interest.

This work raises the possibility that at least for some taxa and regions, anthropogenic land use has significantly altered the relationship between the compositions of fossil pollen assemblages and the vegetation communities that produced them. This finding does not reduce the utility of fossil pollen data as a paleoecological and paleoenvironmental proxy, but it does indicate that pollen-based inferences about the past may be influenced by recent human activities.

A pessimistic view would note that in many areas, where the history of agriculture is longer and its spread more gradual, disentangling the role of human and climatic effects on palynological records is correspondingly more complex (Li et al. 2014, Marquer et al. 2014). Hence, in this perspective, the consequences of the Anthropocene are not just an uncertain future, caused by rapid change, but increased uncertainty about the past, as the human influence on vegetation composition and function is increasingly difficult to disentangle and vegetation and environments around us increasingly shift away from putatively stable pre-historic baselines. Even the pre-settlement era dataset developed here does not wholly remove the signal of human land use as Native Americans were of course active in the prior millennia (Muñoz et al. 2014) but reduces it by focusing on a time period prior to the onset of intensive Euro-American agriculture and timbering.

A more optimistic (and, in our opinion, better-justified) perspective is to note that we can learn much by observing and studying systems that are in flux. One key need is to develop more contemporary and historical calibration datasets from more time periods such as the pre-settlement era data presented here. Another need is to continue to collect and preserve coretop and other “modern” samples and to more precisely parse the samples in “modern” calibration datasets to reflect their actual date of sampling and thereby better link these to the environmental conditions at the time of sampling. Ultimately, we may be moving to a phase of “paleoecology in the Anthropocene” in which we explicitly recognize that even our “modern” calibration samples are being collected from a rapidly changing system. Based on this understanding, we can build calibration datasets (including both fossil samples and their associated environmental characteristics) that comprise data distributed in space and time, and across an ever-increasing range of environmental and anthropogenic conditions. By developing these calibration datasets and analytical tools we gain more power to understand the fundamental processes governing past vegetation dynamics and those governing palynological signals of these dynamics.

## Conclusions

This paper presents a new dataset of pre-settlement era pollen assemblages, furthering understanding of the stability of pollen-vegetation relationships during periods of major land use and vegetation change. We document that for some genera, the relationships between relative pollen proportions and the relative proportions of the parent tree genera have changed, largely as a result of anthropogenic land use. Usage of modern pollen-vegetation relationships to reconstruct past forest composition from pollen data inaccurately predicts the relative abundance of some genera (*Fagus, Betula, Quercus, Pinus, Tsuga*, and *Picea*) on the landscape. These differences are large enough to substantively alter reconstructions of past vegetation from fossil pollen data and necessitate a need for careful consideration of pollen-based estimates of past vegetation.

These results point a new way forward, in which we can build a new generation of multitemporal pollen-vegetation (and pollen-climate) calibration datasets with pollen samples distributed across space and time. By including samples from the pre-settlement era, the late 20th-century calibration datasets, and new surface samples, we have access to increasingly rich calibration datasets. This work supports the use of historical data in reconstructing vegetation from pollen records (Dawson et al. in revision., Jacobson and Grimm 1986, Hotchkiss et al. 2007, Paciorek and McLachlan 2009), and points to the need for more research to disentangle the effects of climate change and land use on past vegetation dynamics (Li et al. 2014). As we bring more data to bear on understanding the distributions and drivers of historical vegetation dynamics through space and time, we can achieve increasingly powerful insights into past vegetation and its relationships to climatic and anthropogenic change. This work helps us assess potential uncertainties in past reconstructions, and provides a way forward for researchers working in areas where historical data are available. In addition, it indicates the need to recognize major anthropogenic land use not only as a process that affects modern forest composition, but as something that can affect our ability to perceive the past.

---

**Supplementary Table 1.**
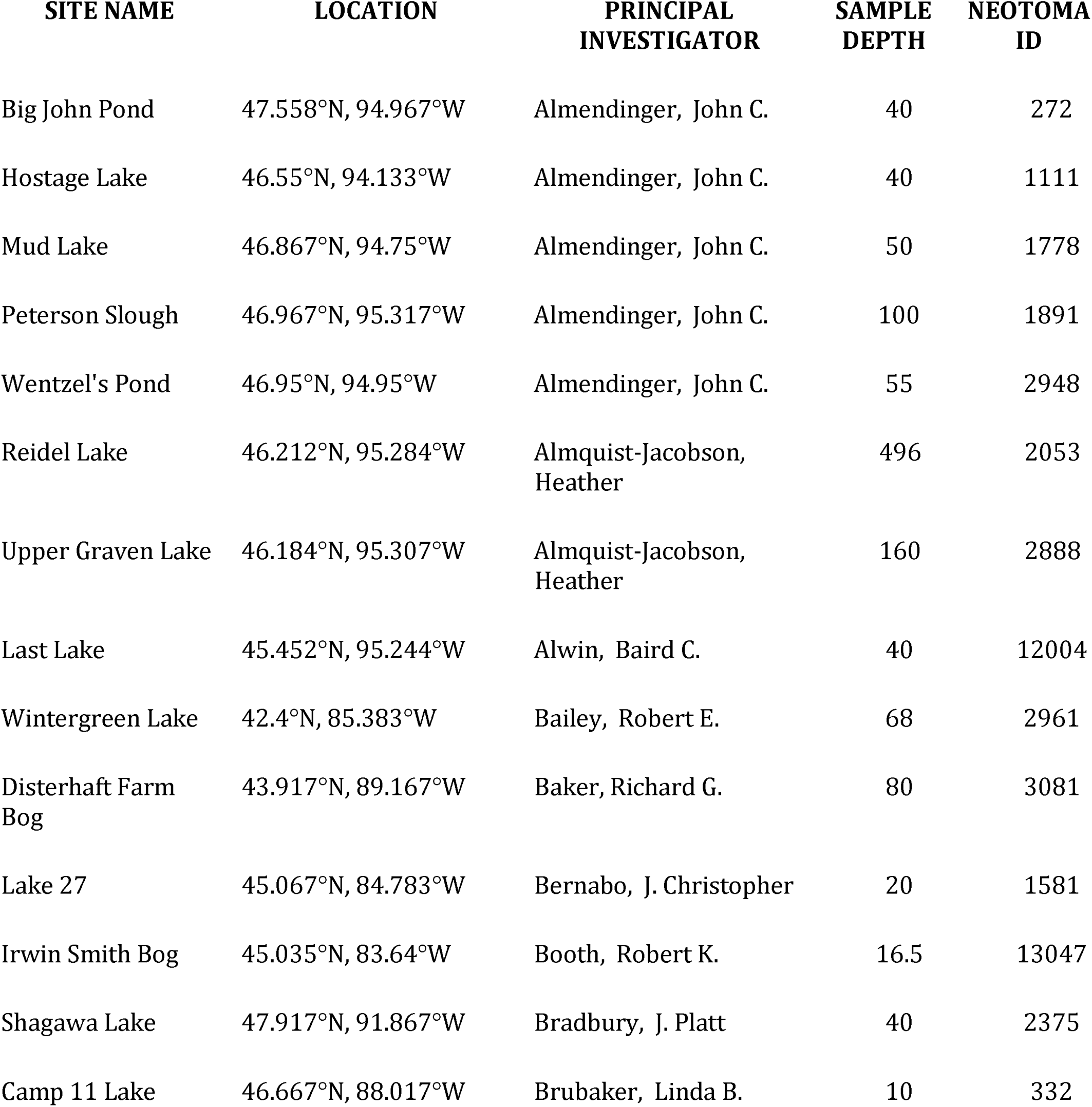
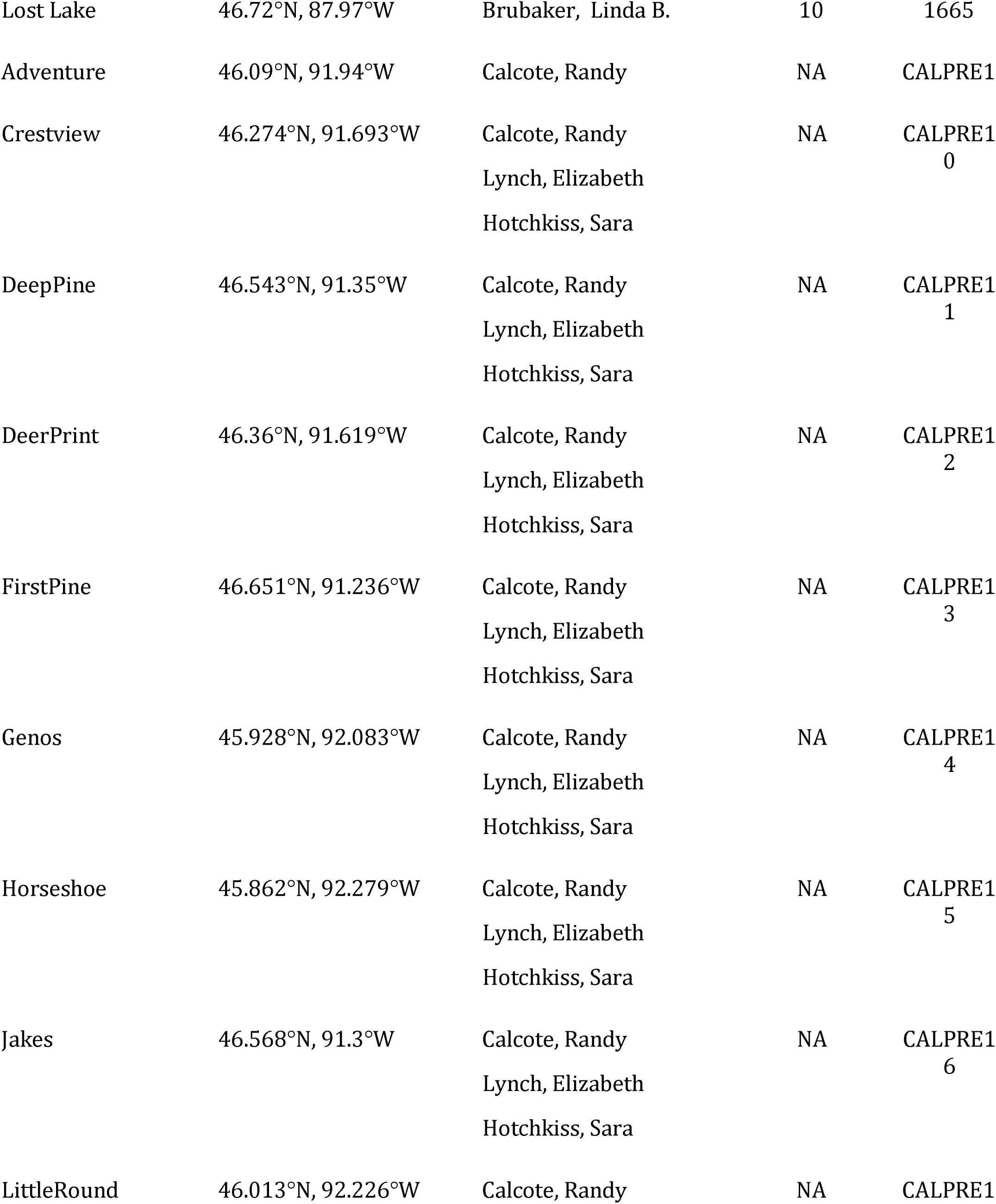
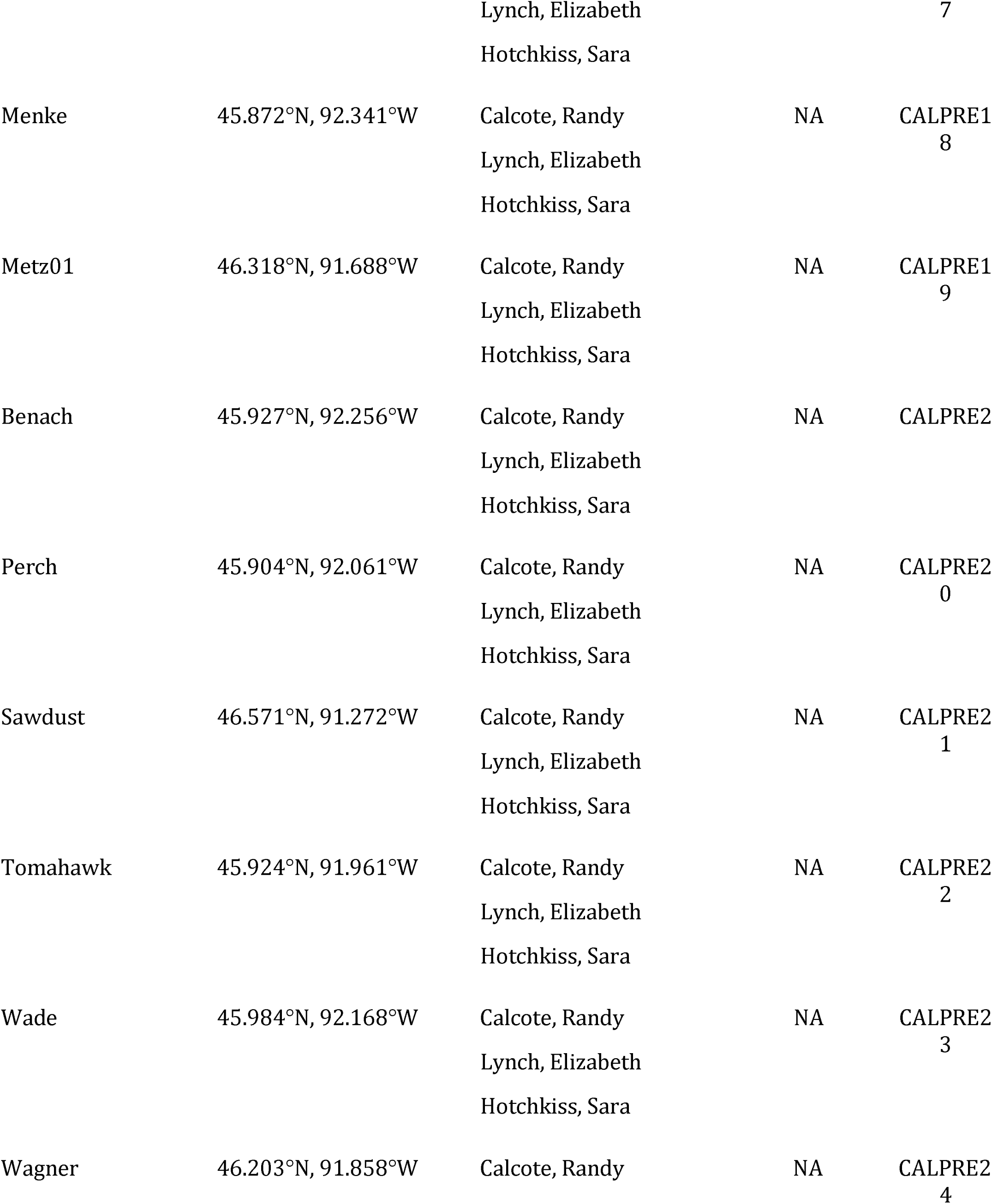
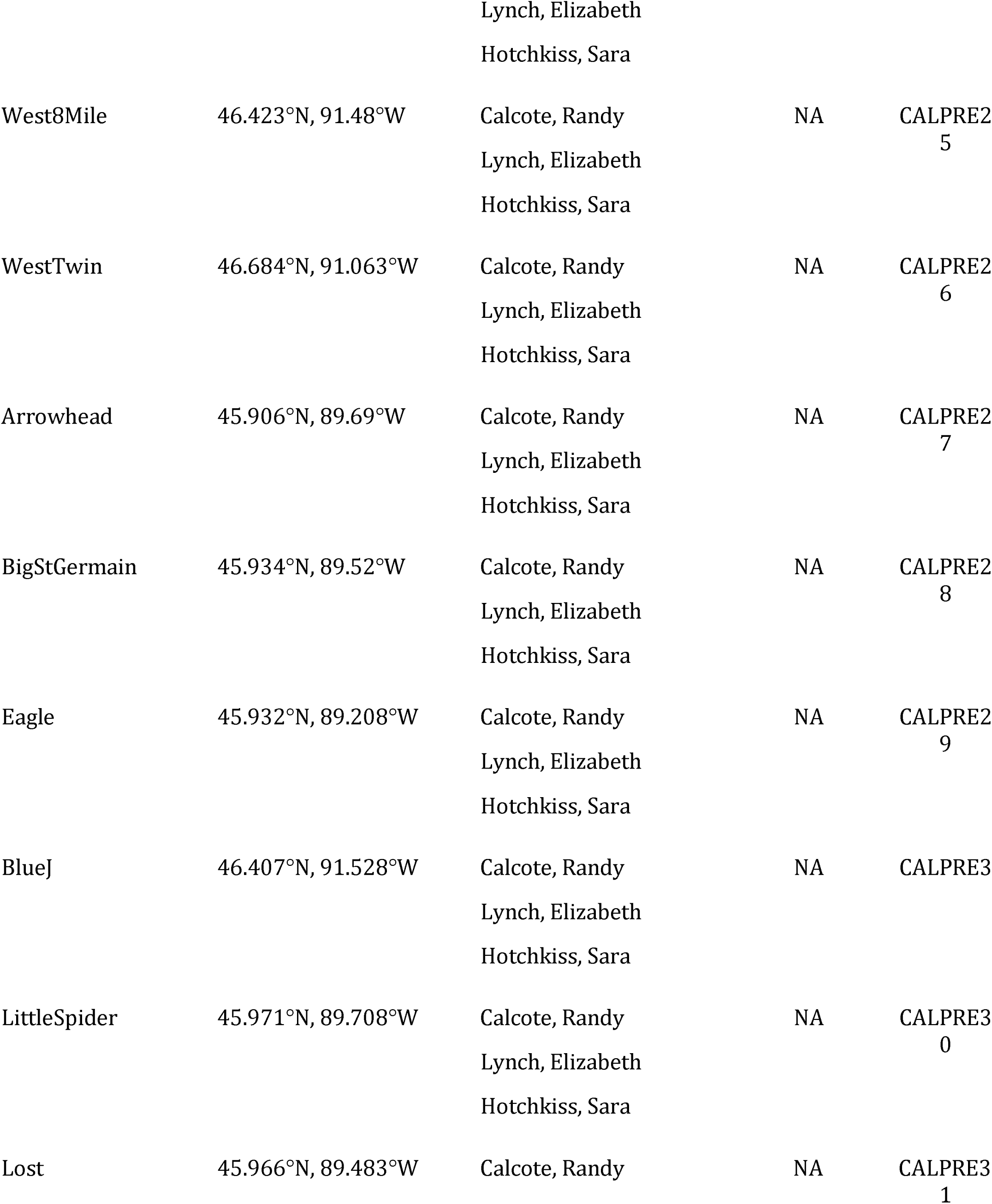
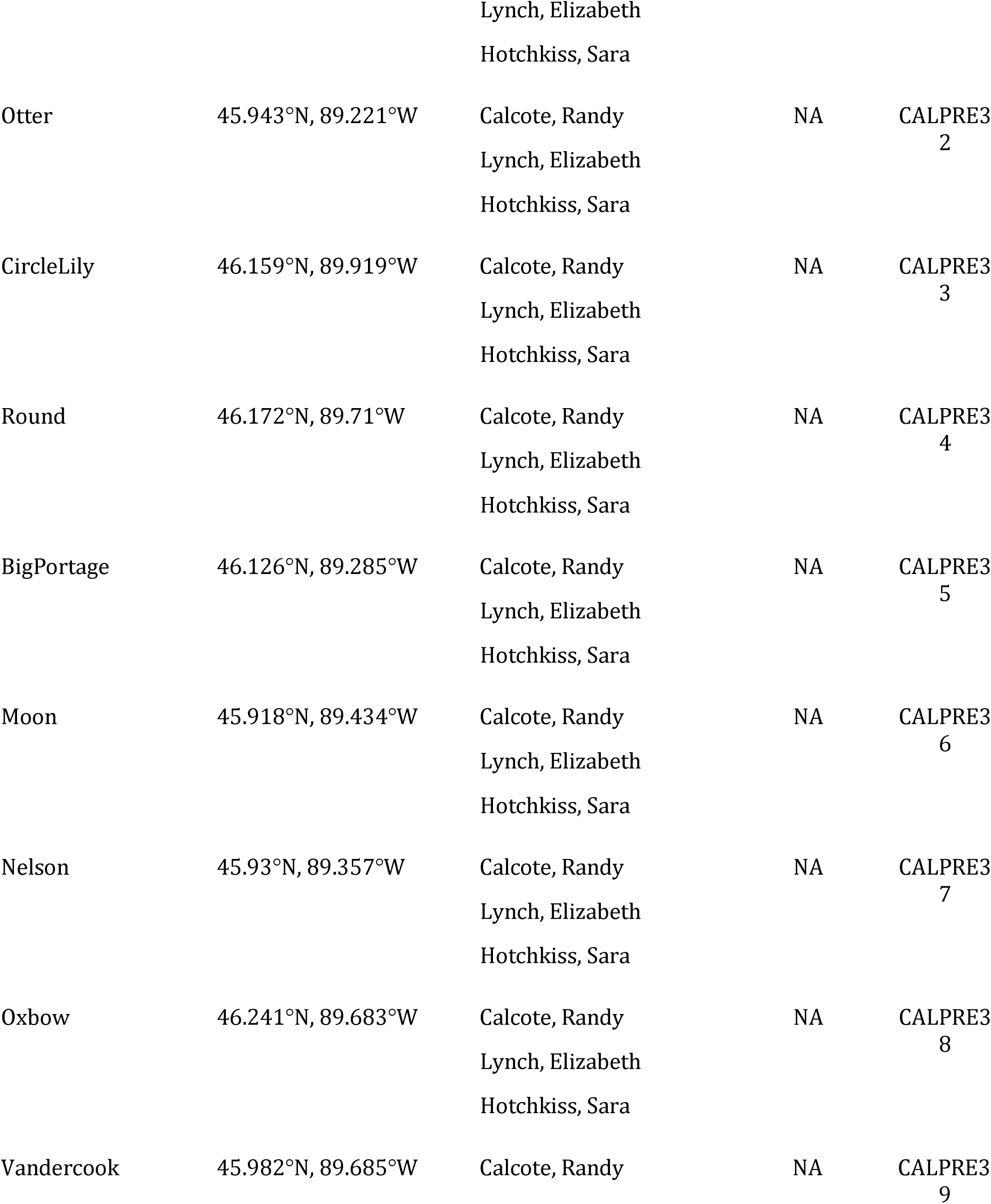
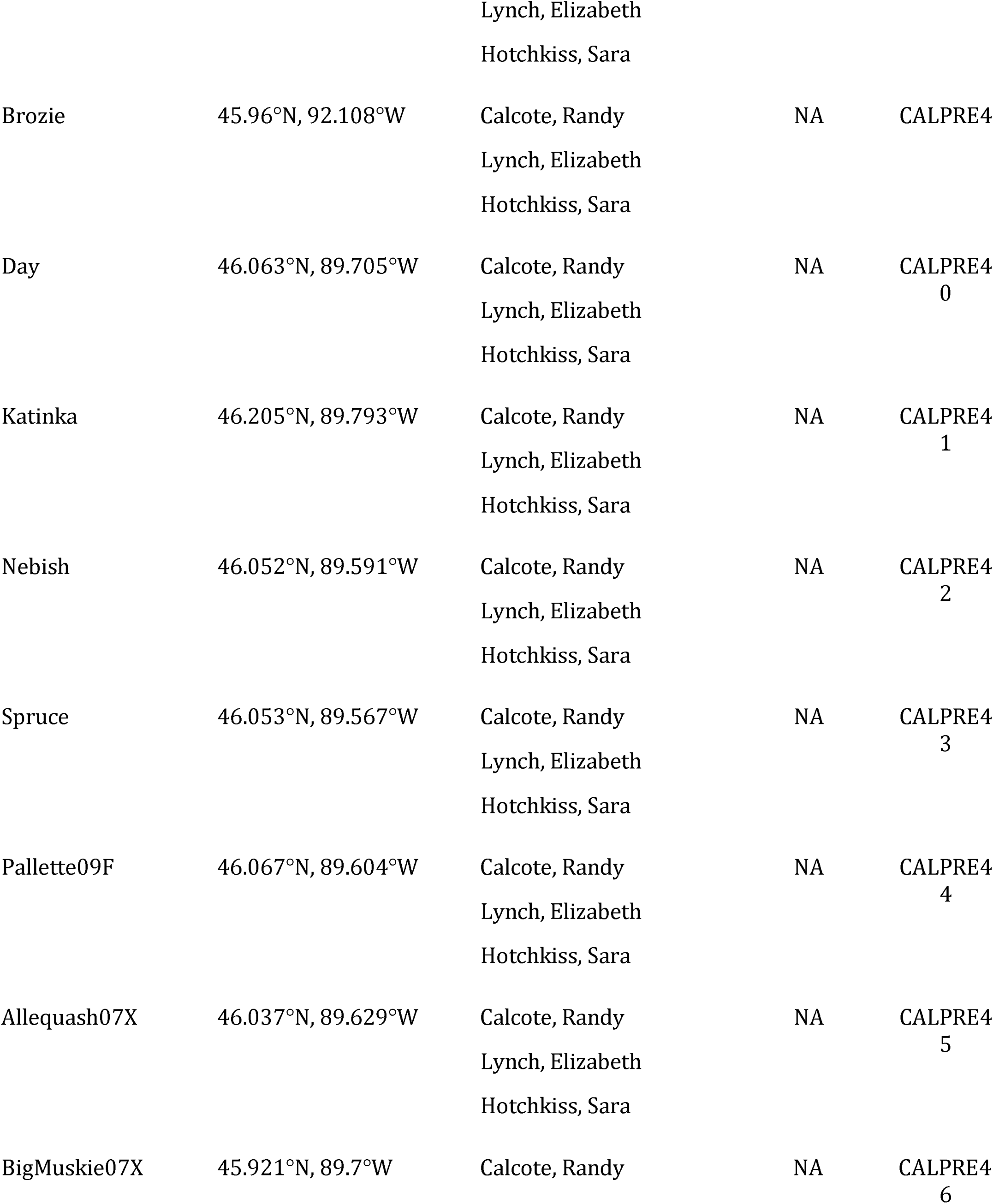
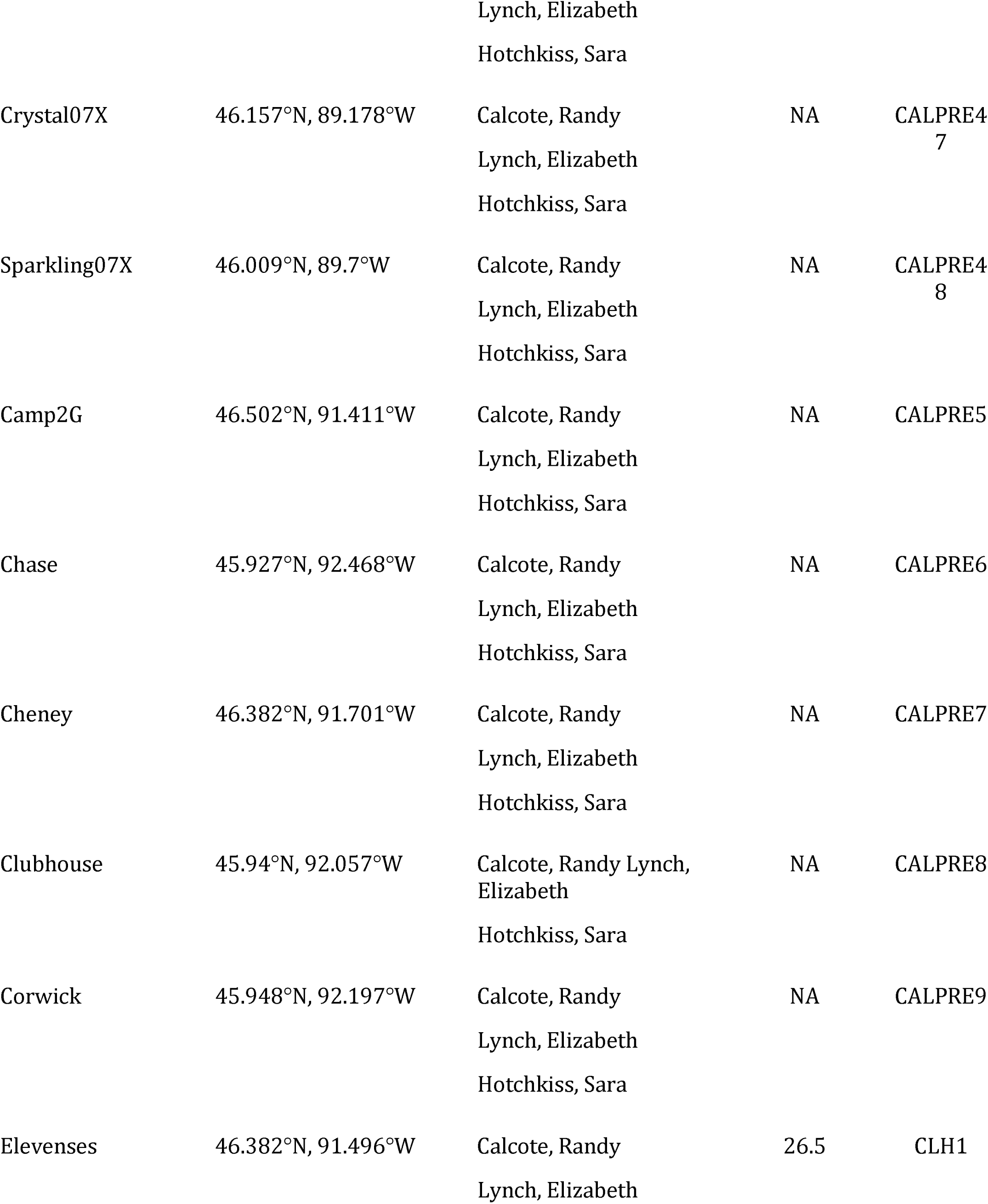
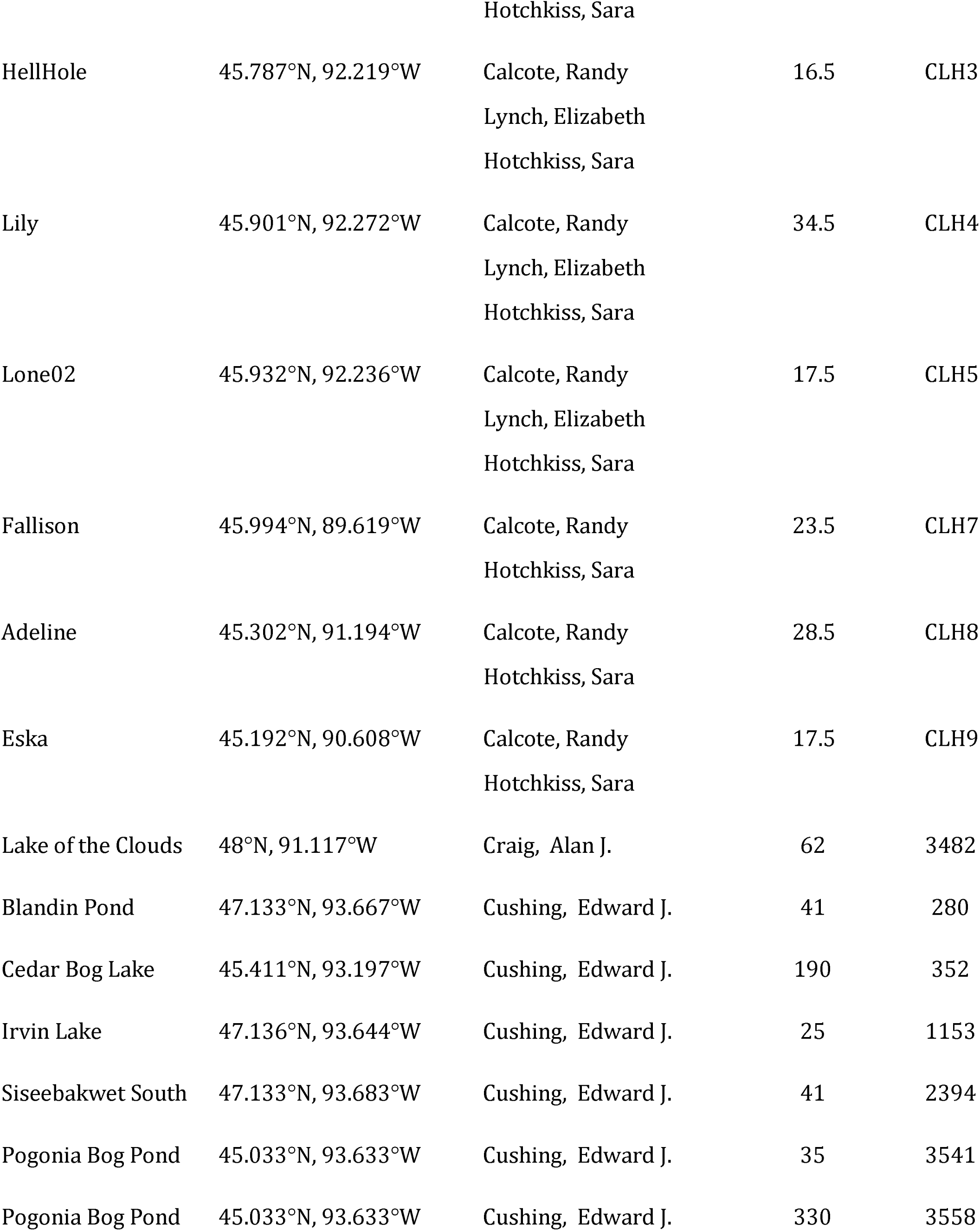
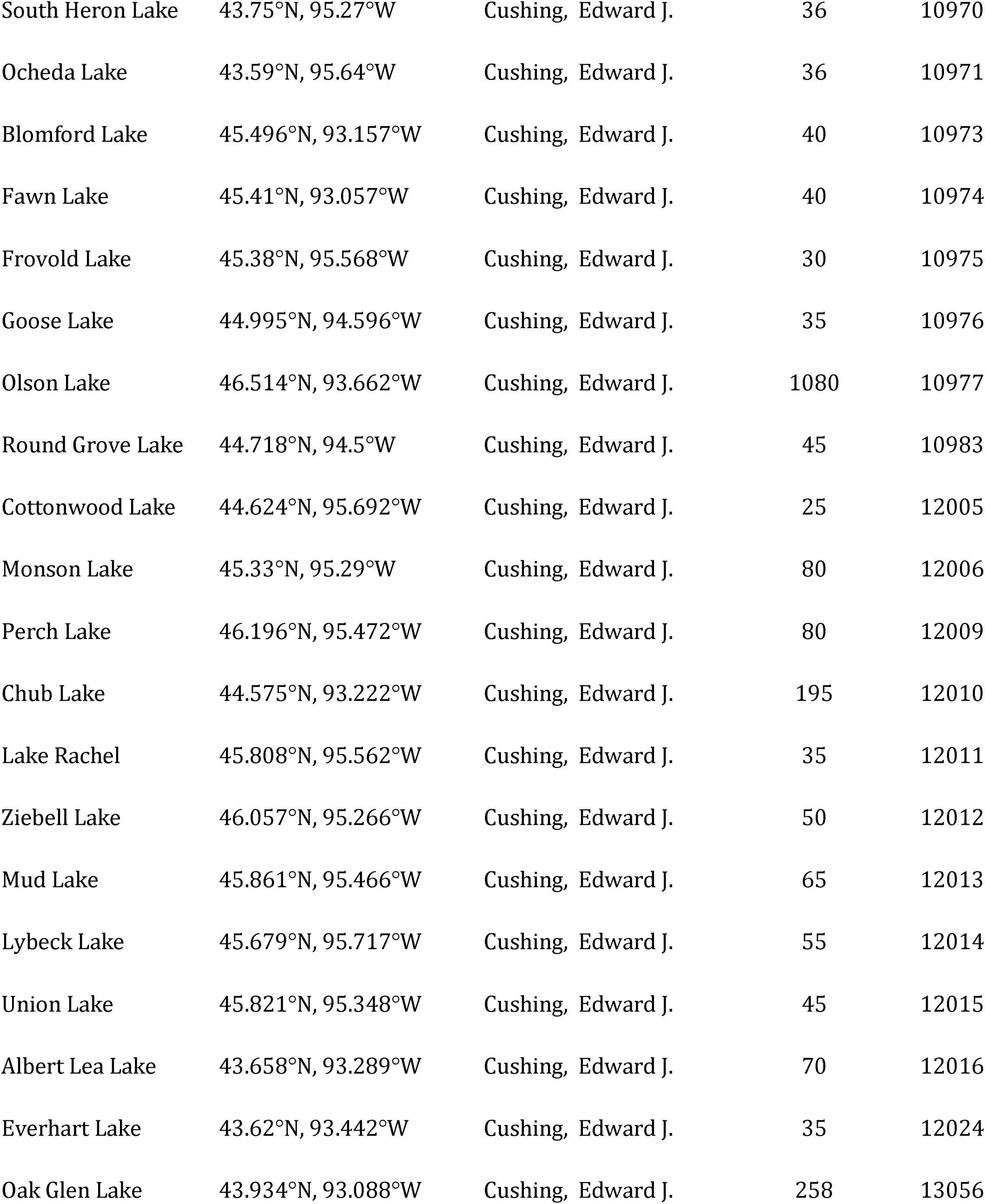
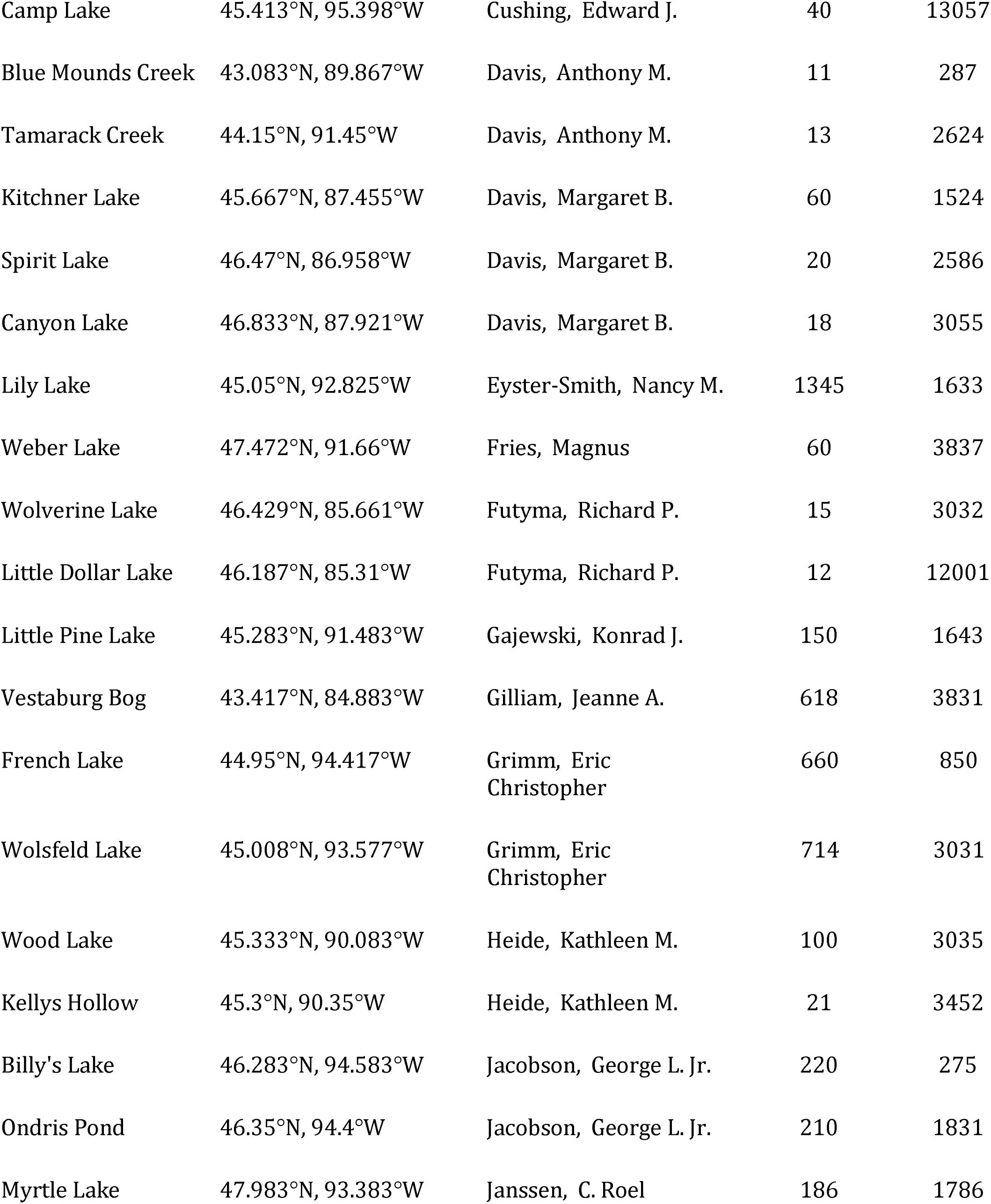
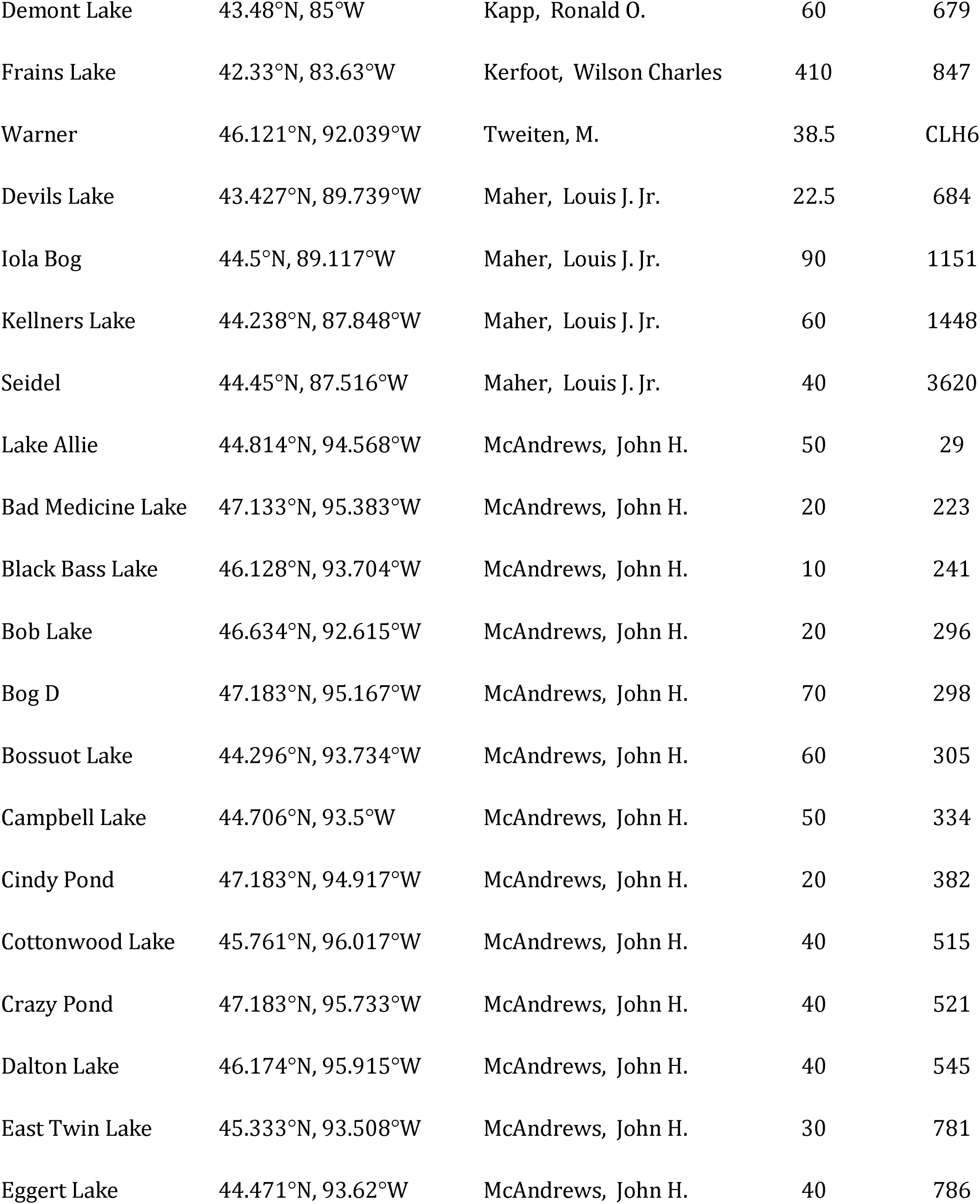
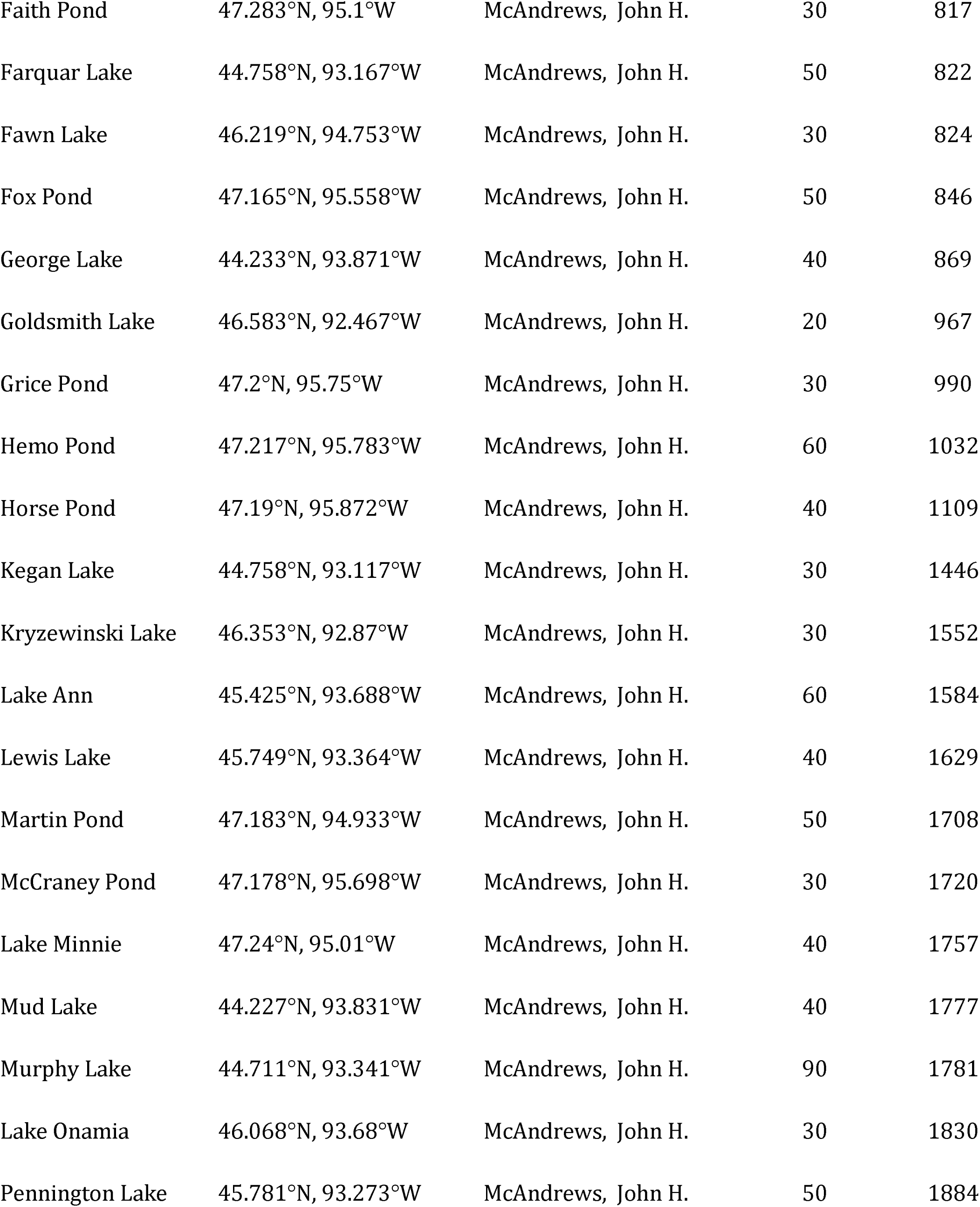
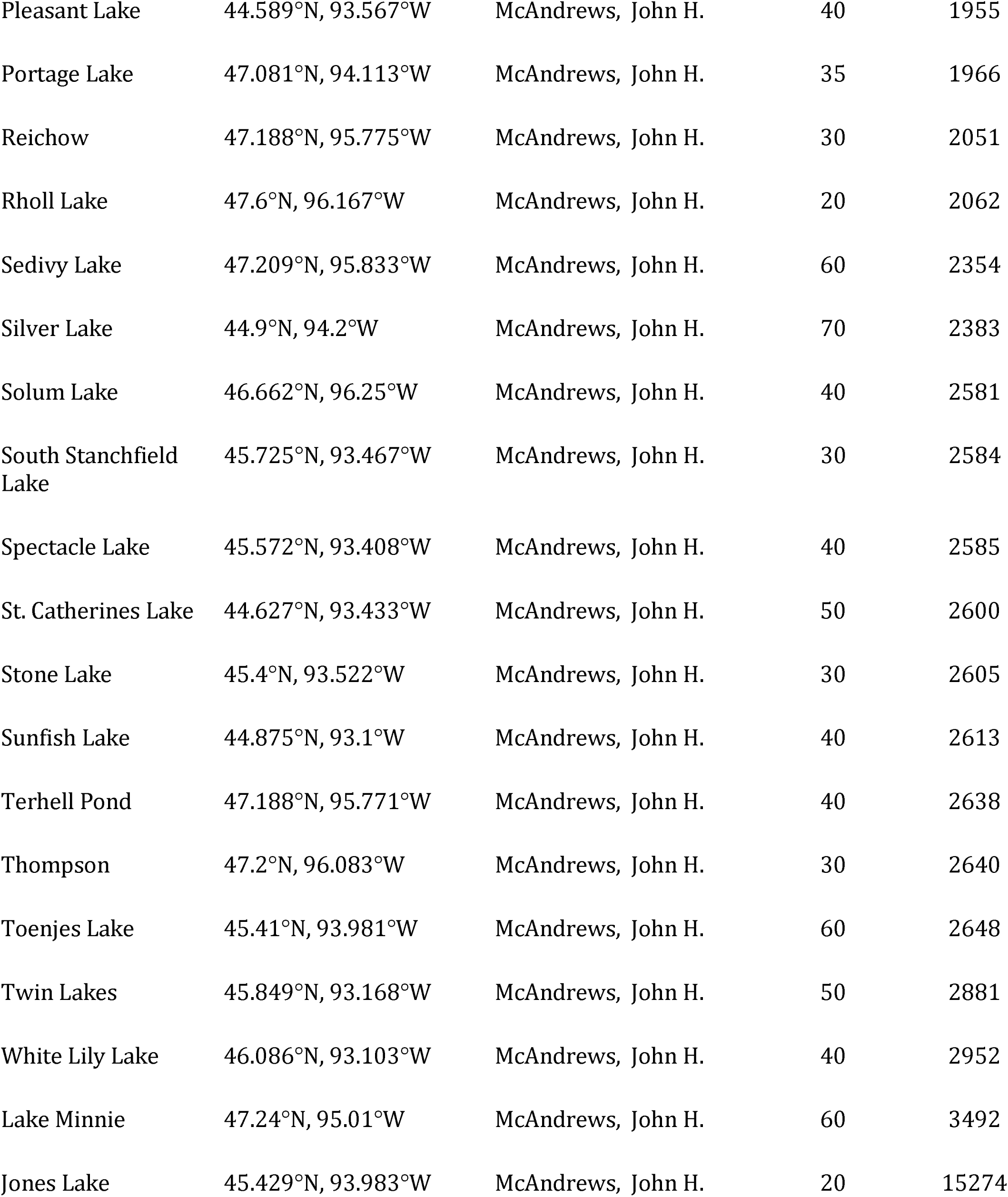
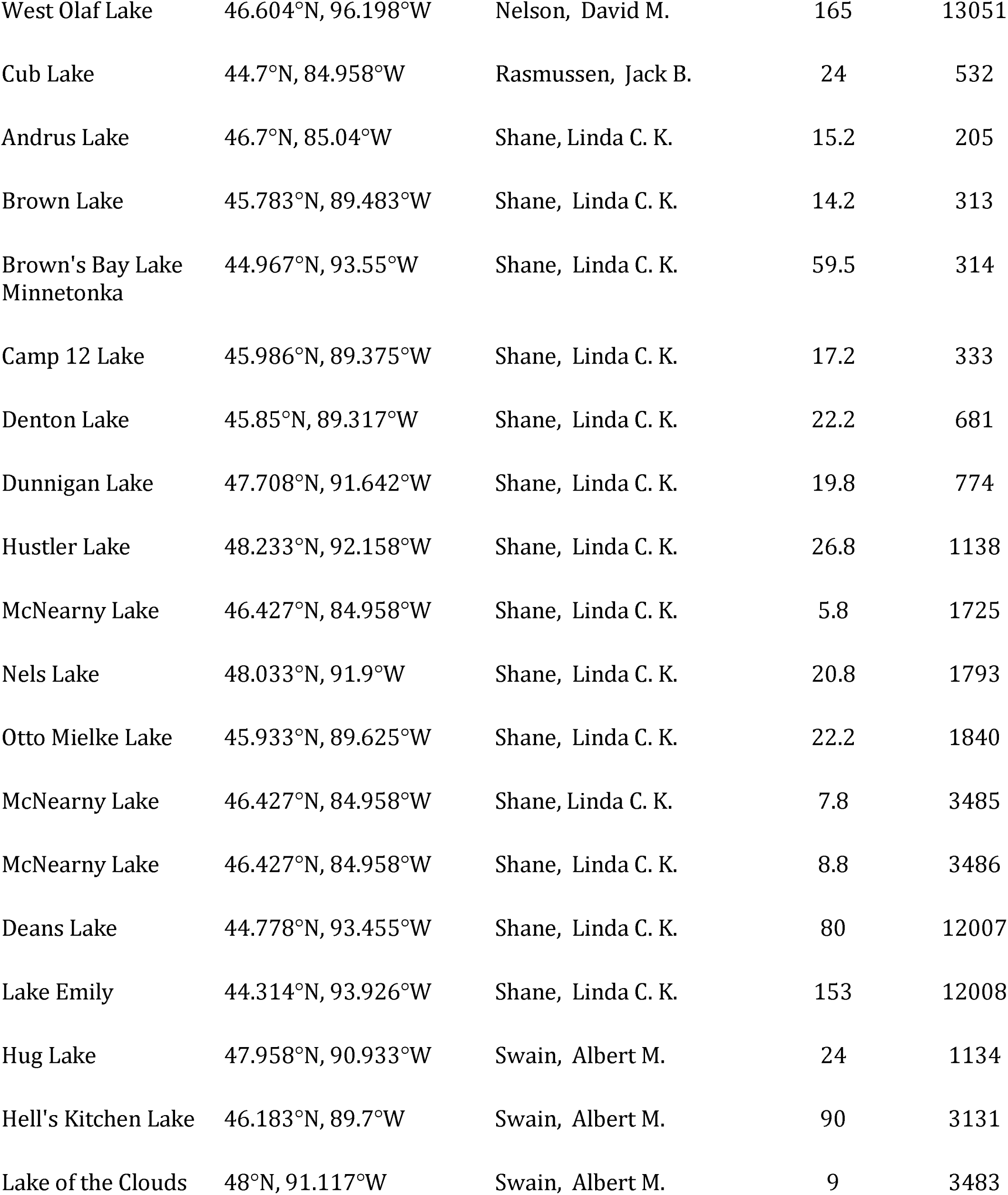
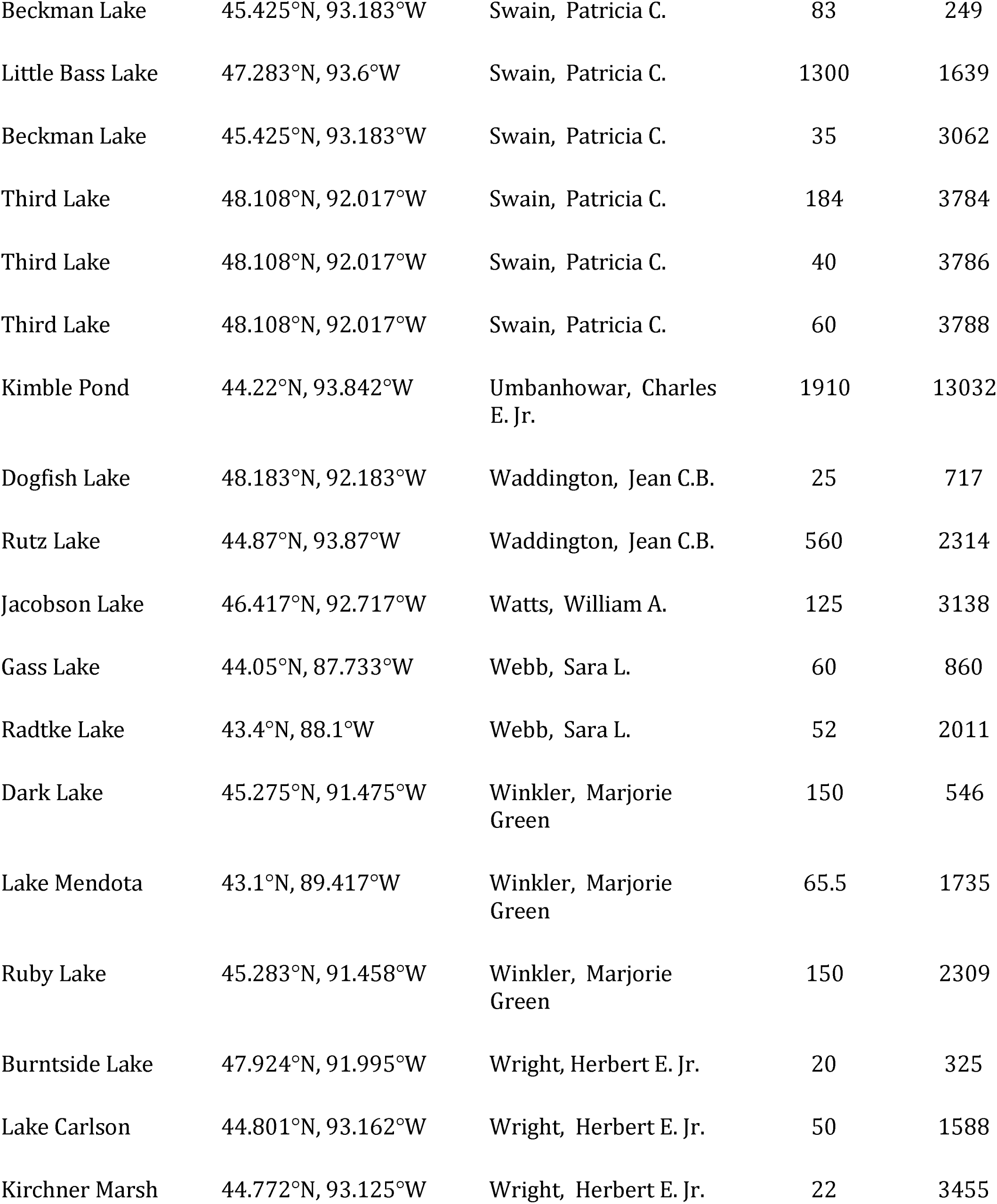
Pollen sample data for samples identified as "pre-settlement" for this study. Table includes the site name, location and PI. The ID number indicates the dataset number within the Neotoma Paleoecological Database; ID numbers that begin with CLH indicate that the sample is from the Calcote dataset, and ID numbers that begin with CALPRE come from the Calcote dataset with only a modern and settlement sample.

